# kinGEMs: A Robust and Scalable Framework for Resource-Constraint Models through Stochastic Tuning of Deep Learning-Predicted Kinetic Parameters

**DOI:** 10.64898/2026.03.14.711833

**Authors:** Rana A. Barghout, Lya Chiñas Serrano, Benjamin Sanchez-Lengeling, Radhakrishnan Mahadevan

## Abstract

The construction of accurate enzyme-constrained genome-scale models (ecGEMs) remains a critical challenge in systems biology, limited by sparse kinetic data and the need for biologically meaningful representations. This work presents an integrated framework combining CPI-Pred, a deep learning model to predict kinetic parameters (***k_cat_***) from sequence and compound embeddings, with kinGEMs, a pipeline to incorporate these uncertainty-aware parameters into ecGEMs for metabolic optimization. By leveraging representations at multiple scales, the approach captures sequence, structure, and kinetic data to enhance model generalizability and accuracy. Rigorous benchmarking demonstrates the framework’s capability to predict growth rates and fluxes that are consistent with experimental observations and measured proteome fractions, reduce median flux variability by several folds, and yield better-constrained metabolic models by explicitly accounting for the uncertainty inherent in ML-predicted kinetic parameters. kinGEMs successfully generated ecGEMs for 93 models spanning a phylogenetically diverse set of organisms including Gram-negative and Gram-positive bacteria, mycobacteria, parasitic protists, fungi, and mammalian cell lines. These innovations breakdown the barrier to application of ecGEMs to non-model organisms and open new avenues for metabolic engineering and synthetic biology in more industrially relevant hosts.

**Author Summary:** Living cells are chemical factories, and computer models of their metabolism can predict how they grow, what they consume, and what they produce, useful for everything from engineering biofuel-producing microbes to understanding disease. The more realistic versions of these models require knowing how fast each cellular enzyme works, but this information is missing for the vast majority of enzymes, especially outside of a handful of well-studied organisms like *E. coli*. We built kinGEMs, an automated pipeline that fills these gaps using a machine learning model to predict enzyme speeds directly from protein sequence, then automatically fine-tunes those predictions so the resulting model’s growth matches real experimental measurements. We show that models built this way better match independent, real-world data they were never tuned to match, including detailed protein-abundance measurements and a classic metabolic behavior (bacteria switching to fermentation at high growth rates) that conventional models cannot reproduce without manual intervention. We also show the approach works reasonably well across 93 different organisms and successfully reconstructs known metabolic trade-offs in an oil-producing yeast unrelated to *E. coli*. This provides a scalable way to build usable, enzyme-aware metabolic models for organisms that currently lack the detailed experimental data such models normally require.

## 2 Introduction

Genome-scale metabolic models (GEMs) have played an essential role in understanding cellular metabolism, enabling predictions of growth phenotypes, metabolic fluxes, and responses to genetic and environmental perturbations. These models provide an interpretable mechanistic framework for metabolic engineering and synthetic biology, a critical advantage over black-box virtual cell models. However, traditional stoichiometric GEMs, constrained only by mass balance and thermodynamics through flux balance analysis (FBA) [1–3], often produce solution spaces too broad for actionable predictions. Enzyme-constrained GEMs (ecGEMs) address this limitation by incorporating kinetic parameters and proteome allocation constraints, substantially narrowing the feasible flux space. This enhanced precision has proven valuable for applications ranging from strain design [4, 5] to understanding evolutionary trade-offs in metabolic regulation [6, 7].

Despite their promise, the construction of accurate ecGEMs remains severely limited by the availability of kinetic parameters. Experimentally determined turnover numbers (*k_cat_*), Michaelis-Menten constants (*K_M_* ), and inhibition constants (*K_I_* ) exist for only a small fraction of characterized enzymes, creating a fundamental bottleneck for model development. For example, approximately 89% of *E.coli^′^s* metabolic enzymes (MG1655) are missing *k_cat_* values from BRENDA [8]. Automated tools like AutoPACMEN have been developed to systematically retrieve available *k*_cat_ data from BRENDA and SABIO-RK [8, 9] for GEM reactions, but coverage remains limited even with multi-database integration [10]. This data sparsity is even more amplified when ecGEMs are extended beyond model organisms or incorporate non-native pathways in metabolic engineering applications. Recent advances in machine learning offer promising solutions to this challenge. Transformer-based protein language models such as ESM [11] and ProtTrans [12] have demonstrated a remarkable ability to extract meaningful sequence representations for downstream prediction tasks, while molecular fingerprints derived from d-MPNN [13] and ECFP [14] frameworks enable prediction of compound-protein interactions. Related approaches have shown that combining quantum chemistry or structural descriptors with machine learning can accurately predict biochemical properties of large reaction spaces [15]. Building on these foundations, several recent methods have applied deep learning to predict enzyme kinetics from protein sequence data and compound information, including DLKcat [16], UniKP [17], RealKcat [18], and CatPred [19], which vary in their underlying architectures (e.g., pretrained protein and molecule embeddings, deep ensemble or Bayesian uncertainty estimation) and in the scope of kinetic parameters predicted. Earlier work on regulatory and context-specific FBA has demonstrated how integrating transcriptomics and regulatory logic with GEMs can refine feasible fluxes and phenotypes, yet these methods typically lack explicit enzyme kinetics [20]. Subsequent frameworks such as GECKO [21] and sMOMENT [10] addressed this gap by advancing the integration of kinetic constraints into GEMs, demonstrating improved predictions of growth phenotypes.

However, critical gaps still remain in translating kinetic parameter predictions into functional, scalable ecGEMs. First, existing benchmarks have focused primarily on the accuracy of individual *in vivo* kinetic parameter predictions in isolation (using correlation metrics), without systematically evaluating how prediction errors propagate to model-level phenotypes such as growth rates and flux distributions under perturbations [19, 22, 23]. Second, current approaches generate static parameter estimates that fail to capture the dynamic nature of enzyme kinetics, which vary with environmental conditions, regulatory states, and metabolic contexts. Cells continuously respond to environmental perturbations through proteomic responses in the form of allosteric regulation, post-translational modifications, and substrate/product availability, mechanisms that fixed enzyme kinetic values typically derived from *in vitro* assays cannot represent [24]. This limitation is particularly problematic when models encounter unrealistic enzyme constraints that produce infeasible solutions [21], limiting both scalability and biological realism. To address this, most approaches set unrealistic proteome mass fractions that overestimate the amount of available active enzymes available in the cell for metabolism [10, 21]. Hence, malleable kinetic parameters constraints are required. Third, the generalizability of kinetic parameter prediction methods across taxonomically diverse organisms has not yet been demonstrated at scale. Existing approaches such as DLKcat [16] have been trained on large corpus of organism data but validated predominantly in single kingdom (Fungi), and have not been systematically applied to phylogenetically distant species where enzyme functional annotation is sparse. More recent predictors such as UniKP [17] and RealKcat [18], while trained on similarly broad, multi-organism datasets curated from BRENDA [8] and SABIO-RK [9], have to our knowledge not been demonstrated as inputs to ecGEM construction. As a result, the construction of ecGEMs has remained limited to a narrow set of organisms, leaving the broad application of enzyme-constrained modeling to less-studied but industrially and clinically relevant microorganisms largely unexplored.

To address these gaps, we developed a comprehensive pipeline that couples kinetic parameter prediction with automated ecGEM construction, optimization, and validation (Fig 1). We utilize CPI-Pred (**C**ompound-**P**rotein **I**nteraction **Pred**iction), a multi-modal deep learning model for kinetic parameter prediction [25] that combines protein language model embeddings with molecular encodings to predict *k_cat_*, *K_M_* , *K_I_* , and *k_cat_/K_M_* (Fig 2A). These predictions are then automatically integrated into a baseline GEM to generate an ecGEM with enzyme capacity constraints formulated to handle isoenzymes, enzyme complexes, and promiscuous enzymes (Fig 2B). To assess biological plausibility, the resulting ecGEM is benchmarked against experimental measurements under different conditions (Fig 2C). Finally, because ML-predicted kinetic parameters are imperfect and condition-dependent, kinGEMs employs stochastic tuning (simulated annealing optimization) loop that iteratively refines *k_cat_* values within relative CPI-Pred-derived uncertainty bounds, using organism-specific growth as validation signals to reconcile molecular-level predictions with systems-level phenotypes (Fig 2D).

**Fig 1:**
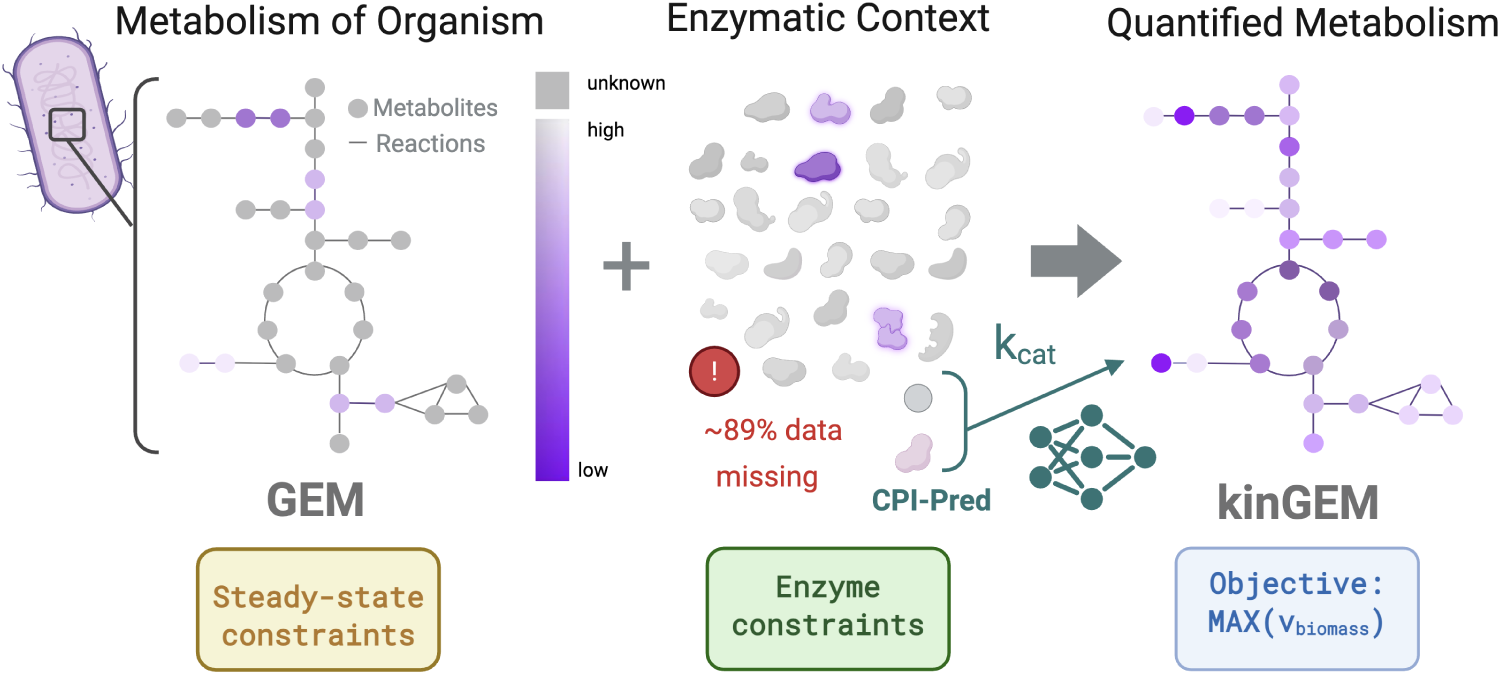
From genome-scale models to quantified metabolism. Overview of the construction of enzyme-constrained genome-scale models (ecGEMs). A conventional genome-scale model (GEM) represents organismal metabolism using stoichiometric reactions subject to steady-state constraints. Incorporating enzyme abundances and catalytic turnover rates (*k_cat_*) introduces enzymatic constraints that couple reaction activity to enzyme capacity, yielding an ecGEM. Despite the improved physiological realism, the application of enzymatic constraints is limited by incomplete kinetic data; for example, approximately 89% of the required *k_cat_* values are missing for *E. coli* models (gray-colored enzymes & reactions). In this work, we address this gap by introducing deep learning–predicted *k_cat_* values and refining them to better align model behavior with biological observations. The resulting kinGEMs framework enables enzyme-constrained quantification of metabolic activity, where reactions are systematically limited by enzyme availability and catalytic efficiency (purple-colored enzymes & reactions).

**Fig 2:**
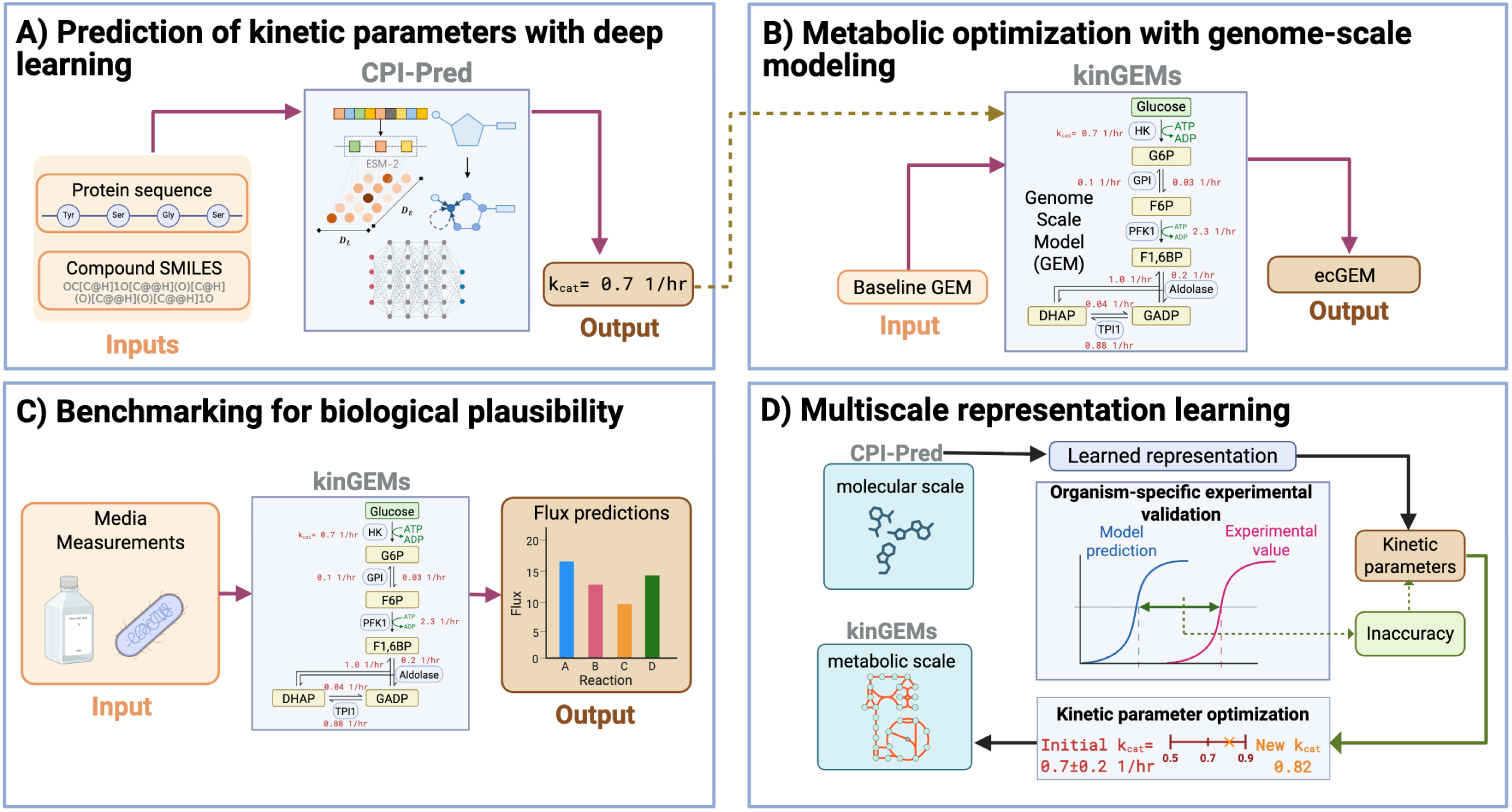
Overview of the integrated CPI-Pred and kinGEMs framework for enzyme-constrained metabolic modeling. **(A)** CPI-Pred architecture for kinetic parameter prediction. Protein sequences and compound SMILES are encoded using ESM-2 embeddings and molecular fingerprints, respectively. The deep learning model predicts kinetic parameters (*k_cat_*, *K_M_* , *K_I_* , and *k_cat_*/*K_M_* ) from these multi-scale representations. **(B)** Integration of predicted kinetic parameters into genome-scale metabolic models. Baseline GEMs are augmented with enzyme constraints using CPI-Pred predictions to generate enzyme-constrained GEMs (ecGEMs). **(C)** Benchmarking framework for biological plausibility. Media conditions are simulated in kinGEMs to predict fluxes across different reactions. Model predictions are systematically evaluated against experimental flux measurements to assess biological alignment. **(D)** Multiscale representation learning and parameter optimization. CPI-Pred learns representations at the molecular scale (compounds and sequences), while kinGEMs operates at the metabolic network scale. Organism-specific validation compares model predictions against experimental values, with discrepancies used to refine kinetic parameters through simulated annealing optimization, creating a feedback loop between molecular predictions and systems-level validation.

In contrast to prior work that evaluates kinetic predictions in isolation, we benchmark ecGEMs on two axes: (i) precision, defined by the contraction of the optimal feasible flux solution space, and (ii) accuracy, defined by agreement with experimental phenotypes and flux measurements. Finally, we also present the scalability results across 93 BiGG GEMs [26]. Together, these components establish a framework that connects machine learning–based kinetic prediction with genome-scale metabolic modeling, enabling systematic evaluation of how enzyme constraints influence model precision, biological realism, and predictive performance. In doing so, kinGEMs provides a practical framework connecting ML-based kinetic prediction and systems-level phenotype validation, building on prior iterative parameter refinement and phenotype-guided optimization previously explored in ecGEM research.

## 3 Results

To determine whether integrating machine learning–predicted kinetic parameters leads to more informative metabolic models, we evaluated the kinGEMs framework along two complementary dimensions: precision and biological accuracy. Precision reflects the degree to which enzyme constraints reduce the feasible metabolic solution space, while accuracy captures agreement between model predictions and experimentally measured phenotypes and intracellular fluxes. We note that this usage is specific to the constraint-based modeling context of this work and differs from conventional statistical usage of these terms; here, precision is a property of the model’s constraints assessed independent of any experimental comparison, while accuracy is assessed only against experimental data. Using the *E. coli* iML1515 model [27] as a benchmark system, we first examine how progressively incorporating enzyme capacity constraints reshapes the feasible flux landscape. We then assess whether these constraints improve alignment with experimentally measured metabolic fluxes and growth phenotypes, and analyze the role of uncertainty-aware parameter tuning in reconciling molecular-scale predictions with system-level behavior.

### 3.1 Precision: Constraining the Solution Space

We utilized flux variability analysis (FVA) to assess how enzymes constraints introduced by the kinGEMs pipeline affected model precision (further described in the Methods section). We performed an ablation study as well as a flux range comparison with experimental metabolic flux analysis (MFA) data to analyze the precision of the kinGEMs models as compared to baseline models.

#### 3.1.1 Progressive constraint integration reduces flux variability

Across all reactions, the introduction of enzyme constraints led to a systematic reduction in flux variability of the near-optimal space compared to the baseline GEM. Fig 3 shows the cumulative distribution of flux variability for reaction *i* across constraint levels. As additional enzymatic constraints are incorporated, the cumulative distributions shifted left, indicating tighter feasible rangers for an increasing fraction of reactions. The progressive addition of constraints also narrows the distributions and shifts them to lower variability (panel B in Fig 3). Results for the non-optimal FVA case, where no optimal biomass is set, can be found in the Supplementary Information Fig A2.

**Fig 3:**
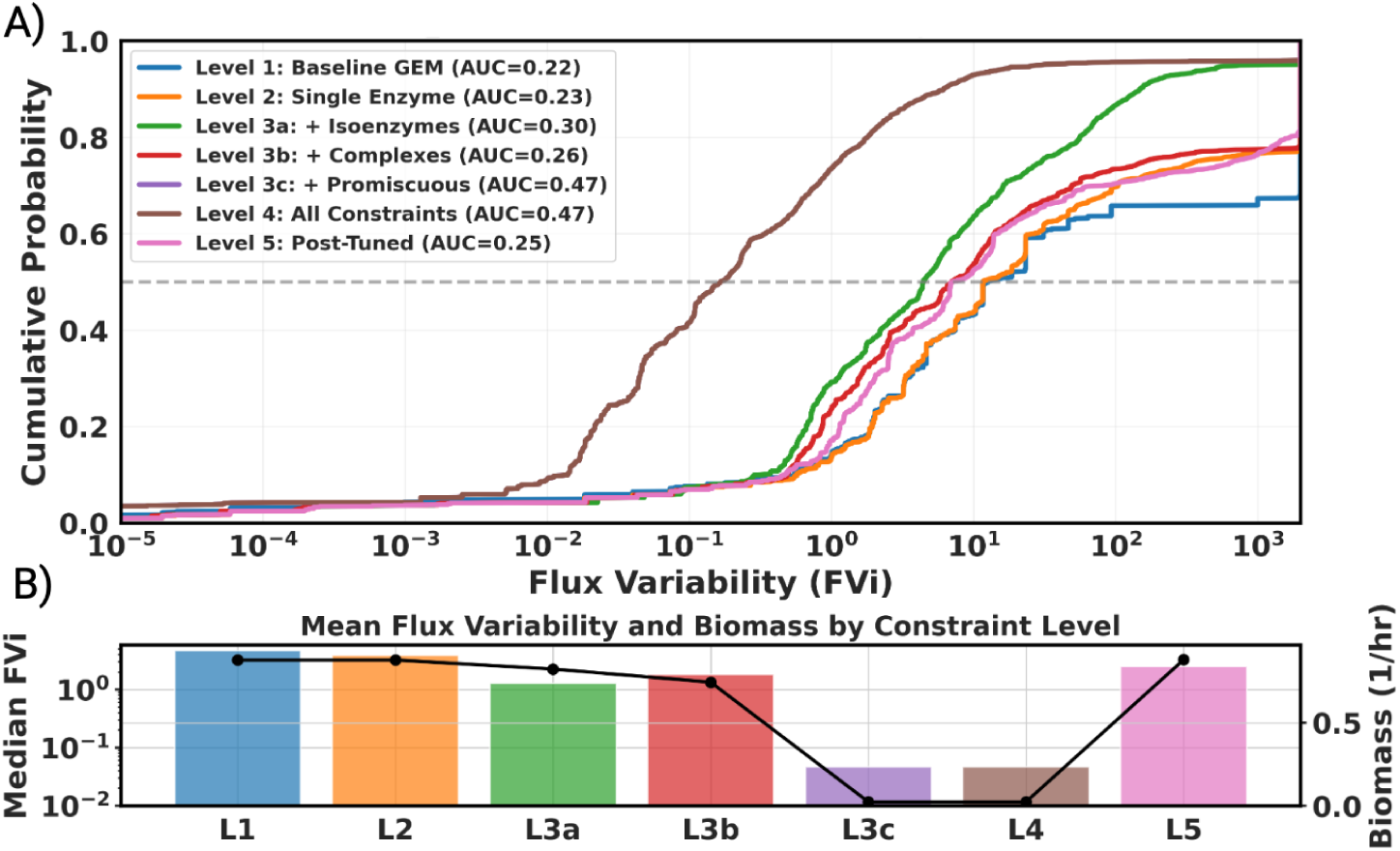
Flux variability ablation study. Progressive integration of enzyme constraints systematically reduces the **optimal** feasible flux solution space. **A)** Cumulative distribution probability of flux variability (FVi) across all reactions for each constraint level. Leftward shifts of the curves indicate tighter feasible flux ranges for an increasing fraction of reactions as enzymatic constraints are added. The dashed line denotes the median FVi. Here, 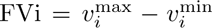 denotes the flux variability of reaction *i*. AUC here denotes area under the curve, where a higher value indicates greater proportion of reactions with low flux variability. *The level 3c line in the plot overlaps with Level 4, and is therefore not clearly visible in the plot. **B)** Median flux variability (bars, left axis) and corresponding optimal biomass flux (points, right axis) for each constraint level. While stronger constraints substantially reduce flux variability, they introduce trade-offs with achievable growth. Post hoc tuning of kinetic parameters (L5) fully restores biomass production while maintaining lower variability than L1.

In particular, the median flux variability decreased monotonically from the baseline GEM (median FVi = 4.7) through intermediate constraint levels, reaching its lowest value under the promiscuous and fully constrained level models with pre-tuned kinetics (median FVi = 0.05). Relative to the baseline GEM, this corresponds to a 94-fold reduction in median variability. However, the growth rate predicted by the ecGEM also drastically decreases in response to these constraints necessitating the additional tuning of kinetic parameters. Although the kinetic parameter tuning increased the median FVi (2.5), we still observe almost a 2-fold reduction compared to the baseline GEM. These results indicate that enzyme-informed constraints substantially restrict the solution space while maintaining optimal growth.

In particular, the addition of promiscuous enzyme constraints (L3c) produced a pronounced drop in median flux variability and growth rate, suggesting that accounting for enzyme reuse across reactions plays a critical role in limiting alternative flux distributions. This result suggests the prevalence of these promiscuous enzymes in key parts of central metabolism.

##### Trade-offs between precision and achievable growth

Although increasing constraint levels consistently reduced flux variability, it also impacted the maximum achievable biomass flux (panel B in Fig 3). Intermediate constraint levels preserved biomass production close to the baseline value of 0.87 1*/hr* for *E. coli* [27, 28], while the fully constrained model exhibited a decrease in growth (97.4% relative to the baseline). Simulated annealing-based tuning alone recovers the substantial majority of this deficit, restoring biomass production to approximately its baseline value while maintaining low flux variability (Supplementary Figures B3). A subsequent growth maintenance parameter (GAM/NGAM) sensitivity sweep (Supplementary Fig B5) shows that any residual growth deficit can be further narrowed by adjusting maintenance-energy assumptions within a biologically bounded range; all results reported here, including in Fig 3, use the literature-default GAM/NGAM values rather than a value selected from this sweep. We also find that running the simulated annealing optimization framework across different seeds produces stable results and converges to the same findings (Supplementary Fig B6).

##### Improved agreement with MFA measurements

To evaluate whether reduced flux variability correlated with improved biological alignment, we compared FVA-derived flux ranges with experimentally measured ^13^C MFA data [29]. Fig 4 panel A illustrates three representative reactions (AKGDH, EX AC e, and CS), chosen as illustrative examples from central carbon metabolism and exchange flux to visually demonstrate that enzyme-constrained models yield flux ranges that are substantially narrower and more aligned with MFA estimates than those obtained from the unconstrained baseline GEM. The full comparison across all 46 mapped reactions is provided in Supplementary Fig C8 and summarized quantitatively in panel B of Fig 4. To assess whether these differences are statistically robust rather than incidental, we performed Wilcoxon signed-rank tests comparing the baseline GEM against kinGEMs pre- and post-tuning models on both mean-to-mean flux distance and FVA interval width (Supplementary Table C2); all comparisons were significant (p *<* 0.001) with large effect sizes (rank-biserial correlation *r* = 0.813–0.929). We note, however, that reduced flux variability (precision) and improved agreement with experimental MFA measurements (accuracy) are related but distinct claims: a narrower feasible flux range does not by itself guarantee closer agreement with the true flux value, since a model could in principle converge on a tight but biologically incorrect range. We therefore rely specifically on the independent mean-to-mean distance comparison against experimental MFA data shown in panel B (rather than the reduction in flux-range width alone) to support our claims of improved biological accuracy.

**Fig 4:**
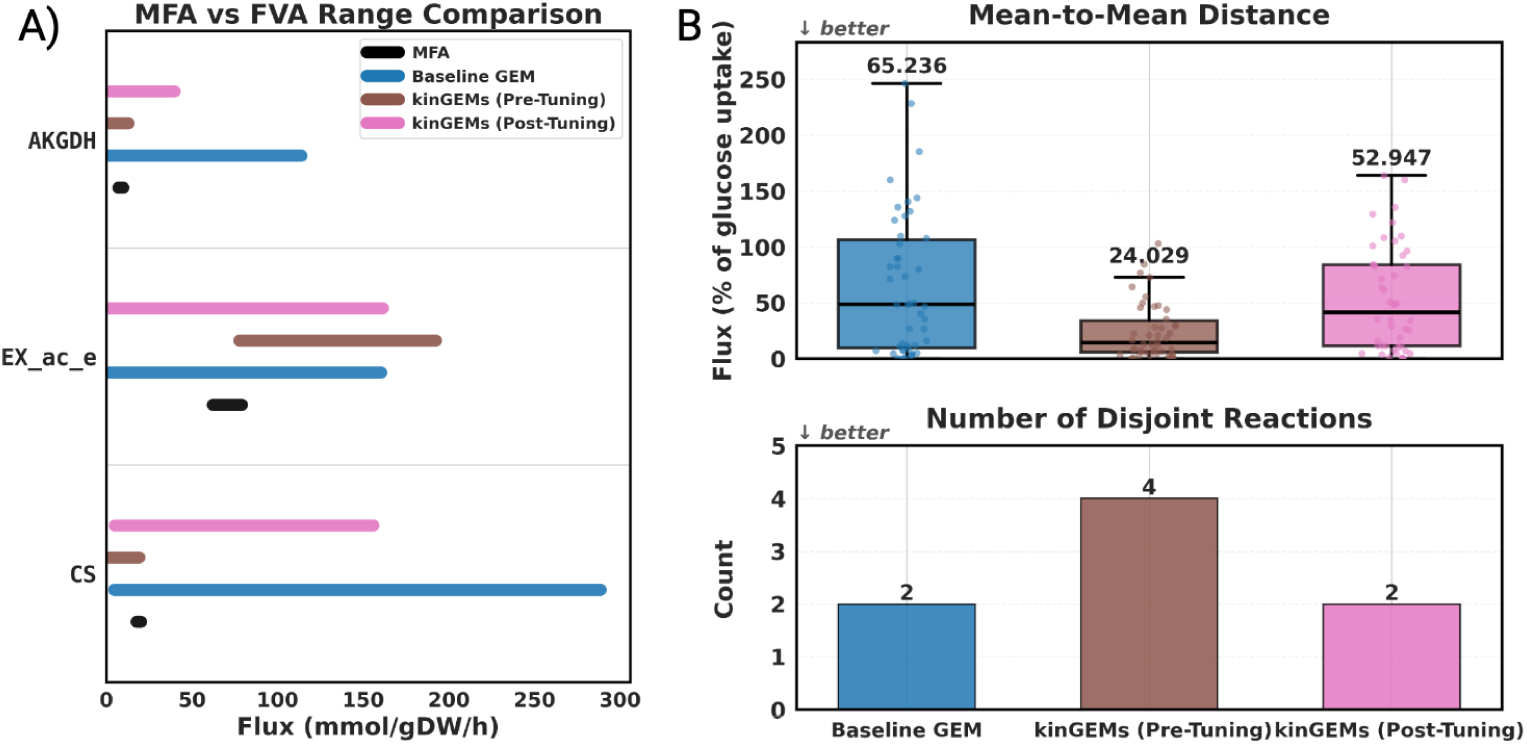
Enzyme constraints improve agreement between feasible flux ranges and experimental MFA measurements. **(A)** Comparison of experimentally measured ^13^C MFA flux ranges (black) with model-predicted feasible flux ranges obtained via flux variability analysis (FVA) for three illustrative reactions: AKGDH (2-oxogluterate dehydrogenase) EX ac e (acetate exchange), CS (citrate synthase). Colored horizontal bars indicate FVA ranges for the baseline GEM (blue), kinGEMs before kinetic tuning (brown), and kinGEMs after kinetic tuning (pink). Enzyme constraints substantially narrow feasible flux ranges relative to the baseline model, bringing predictions closer to experimentally observed values. **(B)** Aggregate agreement across all 46 reactions with available MFA measurements. Top: distribution of mean-to-mean distances in mmol·gDW*^−^*^1^·h*^−^*^1^ between MFA and FVA flux ranges for each model, box-plots summarize distributions, and annotated values indicate the mean-to-mean distance for each model. Bottom: number of reactions with zero overlap between MFA and FVA ranges. While enzyme constraints initially increase the number of disjoint reactions due to tighter feasible ranges, post-tuning of kinetic parameters substantially reduces these inconsistencies and yields the highest overall agreement with experimental flux measurements.

Quantitatively, agreement was assessed using the interval width as well as the mean-to-mean distance between the feasible ranges estimated by the MFA point and those predicted by FVA. Because raw flux magnitudes are directly coupled to each model’s own growth rate (and the pre-tuning ecGEM’s growth rate is itself severely reduced relative to baseline), we rescaled all FVA and MFA flux ranges to each model’s own glucose uptake rate (glucose uptake = 100), following standard ^13^C-MFA convention, before computing these metrics, to avoid conflating a shrunken absolute flux scale with a genuine change in solution-space precision. On this glucose-normalized basis, the pre-tuning enzyme-constrained model’s average FVA interval width was slightly *wider* than the baseline GEM’s (*W*_FVA_ = 9,582 vs. 8,384; Table 1), rather than narrower as suggested by an absolute-unit comparison, reflecting that its apparent absolute-scale tightness was largely an artifact of its collapsed glucose uptake rate (1.37 vs. ∼10 mmol·gDCW*^−^*^1^·hr*^−^*^1^ for the other two models) rather than a genuine contraction of the feasible solution space relative to its own metabolism. Post-tuning, which restores glucose uptake to a comparable scale to the baseline model, achieved the narrowest normalized interval width of the three models (*W*_FVA_ = 7,683), a modest but genuine improvement over baseline. In all cases, FVA interval widths remain substantially broader than the experimental MFA reference (*W*_MFA_ = 3.60 ± 5.03), highlighting that further integration of condition-specific data will be required to approach experimental precision.

**Table 1:**
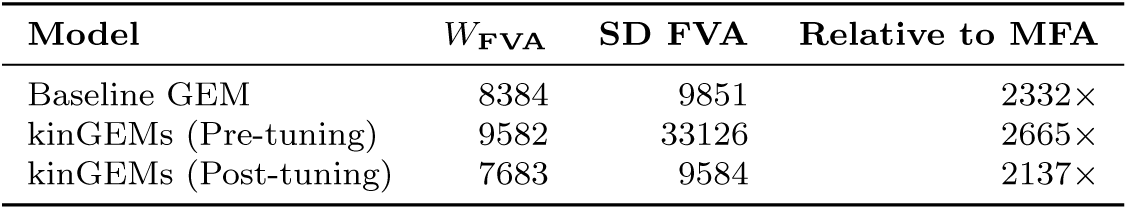
FVA interval width metrics against MFA reference, glucose-normalized (*W*_MFA_ = 3.60 ± 5.03, fluxes rescaled to glucose uptake = 100)

| Model | $W_{\text{FVA}}$ | SD FVA | Relative to MFA |
| --- | --- | --- | --- |
| Baseline GEM | 8384 | 9851 | 2332× |
| kinGEMs (Pre-tuning) | 9582 | 33126 | 2665× |
| kinGEMs (Post-tuning) | 7683 | 9584 | 2137× |

The average mean-to-mean distance, in contrast, preserved the same ranking under normalization: the pre-tuning model showed the closest agreement with MFA (24.0% of glucose uptake), followed by the post-tuning model (53.0%), with the baseline GEM showing the poorest agreement (65.2%; Fig 4, panel B, top). All pairwise comparisons against baseline were statistically significant (Wilcoxon signed-rank test; mean-to-mean distance: *p* ≤ 2.77 × 10*^−^*^4^, *r* = 0.836–0.929; FVA interval width, log scale: *p* ≤ 4.33 × 10*^−^*^4^, *r* = 0.813–0.921; Supplementary Table C2). We note that mean-to-mean distance and interval width capture distinct aspects of model performance and, on the normalized scale, do not agree on which constrained model performs best: the pre-tuning model’s flux distribution shape (relative pathway partitioning) tracks the MFA-observed pattern most closely on average, despite occupying a wider normalized solution space, while the post-tuning model achieves the tightest normalized solution space despite a larger average distance from the MFA point estimate. We discuss the implication of this distinction below. The results of all the reactions analyzed can be found in Supplementary Fig C8.

The impact of parameter tuning is most visible in the analysis of reactions in which FVA failed to overlap with the MFA ranges (Fig 4, panel B, bottom). On the glucose-normalized basis, both the baseline and post-tuning models showed 2 non-overlapping reactions (of 46), while the pre-tuning model showed 4 - a substantial reduction from the 24 non-overlapping reactions apparent under an absolute-unit comparison. This indicates that most of the pre-tuning model’s apparent disagreement with MFA in absolute terms reflects its systematically compressed flux scale, driven by its collapsed glucose uptake rate, rather than a genuinely different flux distribution shape: once this scale offset is removed, the pre-tuning model’s relative flux partitioning was already reasonably well aligned with the experimental data. This reduction in non-overlapping reactions across models was not statistically significant by McNemar’s test (baseline vs. post-tuning: *p* = 1.0; baseline vs. pre-tuning: *p* = 0.625, odds ratio = 0.429; both with *n* = 4 discordant pairs), so we present this comparison descriptively rather than as an additional significance claim. Together, these results indicate that enzyme constraints alone do not substantially distort flux distribution shape relative to experimental measurements, even though the resulting absolute flux magnitudes are severely growth-rate-limited; kinetic parameter tuning’s primary contribution in this analysis is restoring feasible absolute throughput and tightening the normalized solution space, rather than correcting a badly mis-routed flux distribution.

Overall, these results demonstrate that CPI-Pred-derived enzyme constraints, particularly when combined with post hoc kinetic tuning, meaningfully improve agreement with experimental MFA measurements once flux magnitudes are compared on a fair, throughput-normalized basis. As illustrated above, comparing models on an absolute flux scale can be actively misleading when growth rates differ substantially between them, since apparent precision or accuracy gains can simply reflect a compressed flux scale rather than a genuine improvement in the underlying solution space. We therefore report the glucose-normalized comparison as our primary measure of accuracy throughout this study, and emphasize more broadly that flux-range narrowing alone does not by itself establish improved biological accuracy.

### 3.2 Accuracy: Agreement with Proteomic and Phenotypic Data

While the analyses above establish that enzyme constraints and kinetic tuning improve model precision, precision alone does not establish biological realism. We therefore separately evaluated accuracy against quantitative proteomics, an independent data modality not used during kinetic parameter tuning, across 15 growth conditions and 4 independent kinetic parameter sources (CPI-Pred, an independent deep learning predictor CatPred, literature-derived BRENDA values, and randomly sampled log-normal values), using the dataset of Schmidt et al. [30]. Supplementary phenotype-level and generalization analyses supporting this section are provided in Appendix D.

#### 3.2.1 Proteomic agreement across growth conditions and *k_cat_* **sources**

Across all 15 conditions and 4 *k_cat_* sources, predicted enzyme abundances (copies/*gDCW ^−^*^1^, recovered from the model’s own enzyme mass variable) showed consistent, moderate agreement with Schmidt et al. measured protein abundances (Pearson *r* ranging 0.27-0.46 across condition-source combinations, log10 scale). As a complementary error metric, we computed RMSE between predicted enzyme mass and measured proteomics for each condition and *k_cat_* source, both before and after simulated annealing tuning (Fig 5). No single *k_cat_* source dominated this comparison by either metric: CPI-Pred, CatPred, literature-derived BRENDA values, and randomly sampled log-normal values achieved broadly comparable correlation and RMSE with measured proteomics across the majority of conditions tested. Notably, RMSE did not consistently improve after simulated annealing tuning; several conditions show marginally higher post-tuning RMSE than their untuned counterpart (Fig 5), which is consistent with simulated annealing optimizing for growth rate agreement under an enzyme-mass budget, rather than proteomics fit directly.

**Fig 5:**
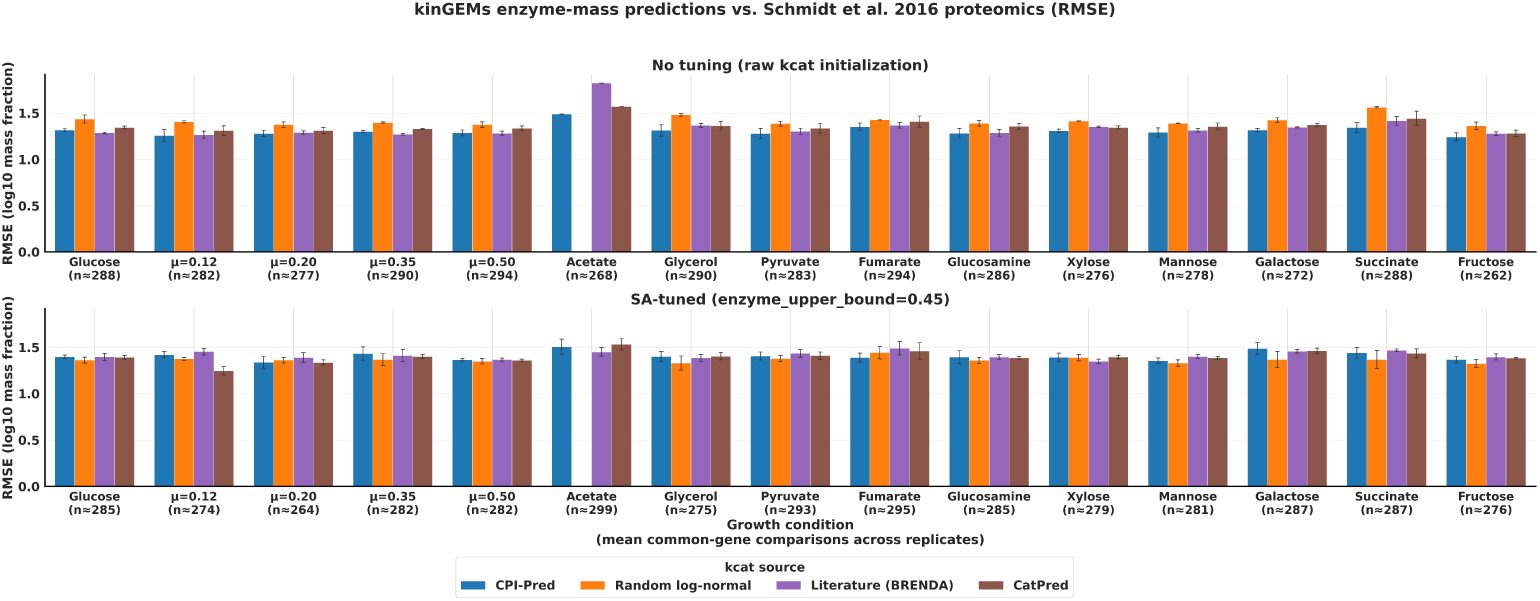
Proteomics agreement (RMSE) across growth conditions and *k_cat_* sources, before and after simulated annealing tuning. RMSE (log10 mass fraction) between kinGEMs-predicted enzyme mass and Schmidt et al. 2016 measured proteomics, across all 15 growth conditions and 4 *k_cat_*initialization sources (CPI-Pred, Random log-normal, Literature/BRENDA, CatPred). **Top:** RMSE using the initial (untuned) *k_cat_* values for each source. **Bottom:** RMSE after simulated annealing tuning (enzyme upper bound = 0.45). Both panels are computed on the replicate-averaged, common-gene basis (the intersection of genes matched across all four *k_cat_* sources for a given condition and replicate), following a single per-condition additive offset that absorbs the fixed per-gDCW-biomass vs. per-cell basis difference between prediction and measurement. Error bars show standard deviation across up to 2 replicates. No single *k_cat_*source is consistently the best- or worst-performing across conditions; the acetate condition shows elevated RMSE relative to other conditions in both panels.

Because single-enzyme *k_cat_* prediction methods (CPI-Pred, BRENDA lookup, Cat-Pred) are not designed to characterize obligate multi-subunit enzyme complexes or translation machinery, we additionally verified proteomics agreement after excluding genes belonging to such categories, identified directly from iML1515’s own gene-protein-reaction rules (271 of 1516 genes, 17.9%, primarily obligate subunits of complexes in oxidative phosphorylation, transport, and outer-membrane porins). This filtering improved correlation modestly and consistently (45 of 58 condition-source pairs improved; mean Δ*r* = +0.018), indicating that out-of-scope genes account for part, but not the majority, of residual prediction error.

#### 3.2.2 Cross-condition generalization of tuned kinetic parameters

A central question for any parameter-tuning procedure is whether the tuned parameters capture generalizable properties of the system or merely overfit to the specific target used during optimization. To test this, we performed a cross-condition generalization analysis: for each of the 15 growth conditions and 4 *k_cat_* sources, we took the *k_cat_* vector produced by simulated annealing under one condition and, without further tuning, re-solved FBA under every other condition’s medium. The resulting predicted enzyme abundances were then scored against that target condition’s own measured proteomics, yielding 767 cross-condition comparisons alongside the 55 matched (native) comparisons.

Mean proteomics correlation was statistically indistinguishable between matched tuning (r = 0.336) and cross-condition transfer (r = 0.335; gap = −0.0006), a pattern that held consistently across all four *k_cat_* sources (per-source gap range: −0.010 to +0.003; full per-condition detail in Fig D11, Appendix D.2). This result is notable because condition identity strongly affects the resulting growth rate itself (predicted growth rate on a fixed medium ranged from *µ* ≈ 0.04 to *µ* ≈ 0.87 h*^−^*^1^ depending on which condition’s tuned *k_cat_* values were applied) yet this large variation in growth outcome did not translate into a corresponding loss of proteomics fit when *k_cat_*s were transferred across conditions. This suggests that simulated annealing predominantly reallocates enzyme mass among a relatively stable set of high-abundance, load-bearing genes largely independent of which specific condition drove the search. We note that a subset of target conditions (acetate, fumarate, pyruvate, galactose) were consistently harder to fit regardless of the source condition or *k_cat_* source used, indicating these reflect intrinsically more difficult proteomics targets rather than source-specific overfitting. As a complementary check restricted to the glucose condition, we also tested whether proteomics-aware tuning of a training gene subset improves fit on a disjoint, held-out gene subset never included in that tuning objective (Methods 5.6.3, Appendix D). Unlike the cross-condition result above, this pathway-level check did not show statistically significant improvement on the held-out gene subset in any completed run (95% bootstrap confidence intervals), indicating that proteomics-aware tuning under this procedure fits the specific pathway it is trained on without demonstrating reliable generalization to a distinct, unseen metabolic pathway within the same condition. This contrasts with the condition-level transfer result above and suggests that whatever generalizes well across growth conditions (likely broad, enzyme reallocation driven by growth rate among high abundance genes) is distinct from what a narrower, gene-subset-targeted proteomics-fitting objective learns.

### 3.3 Generalization to a Non-Model Oleaginous Yeast: *Rhodotorula toruloides* Resource Allocation

The analyses above establish precision, proteomic accuracy, and the recovery of emergent phenotypes within *E. coli*. A more demanding test of generalization is whether kinGEMs recovers *intracellular* resource allocation in an organism it was never tuned against, using only a growth-rate target. We therefore applied the framework to *Rhodotorula toruloides* (*R. toruloides*), an oleaginous yeast whose metabolism (compartmentalized lipid biosynthesis and a non-canonical pentose-assimilation route) is far less kinetically annotated than that of *S. cerevisiae* or *E. coli*. We reconstructed enzyme-constrained models from rhto-GEM [31] for the six carbon-source × growth-phase combinations reported by Rekena et al. [32] (Methods 5.6.4) and compared predicted cofactor and carbon routing against their manually curated GECKO 2.0.2 reconstructions. The two approaches differ sharply in their data requirements: Rekena et al. constrained every reaction with condition-specific per-enzyme abundances and iteratively curated BRENDA *k*_cat_ values, whereas kinGEMs used a single global enzyme-mass pool, CPI-Pred *k*_cat_ predictions, and simulated annealing tuned to the measured growth rate alone. Intracellular fluxes were never part of the kinGEMs objective, so any recovered allocation phenotype is an out-of-sample prediction.

#### 3.3.1 NADPH allocation trade-off between nitrogen assimilation and lipogenesis

Rekena et al. reported that NADPH demand in this oleaginous yeast is partitioned between nitrogen assimilation and lipogenesis: on glucose, NADP-dependent glutamate dehydrogenase (GDH1) carried 46% of NADPH turnover in exponential phase against 13% through fatty acid synthase (FAS1–2), reversing to 12% versus 75% under nitrogen limitation. From a growth-only objective, kinGEMs recovered this switch (Fig 6A): GDH1 fell from 65% to 3% of cytosolic NADPH consumption while FAS1–2 rose from 9% to 88% between the exponential and nitrogen-limited phases. The predicted switch is sharper than the Rekena et al. medians, but its direction (a near-complete reallocation of NADPH from amino-acid synthesis to lipid synthesis under nitrogen limitation) matches the reported trade-off.

**Fig 6:**
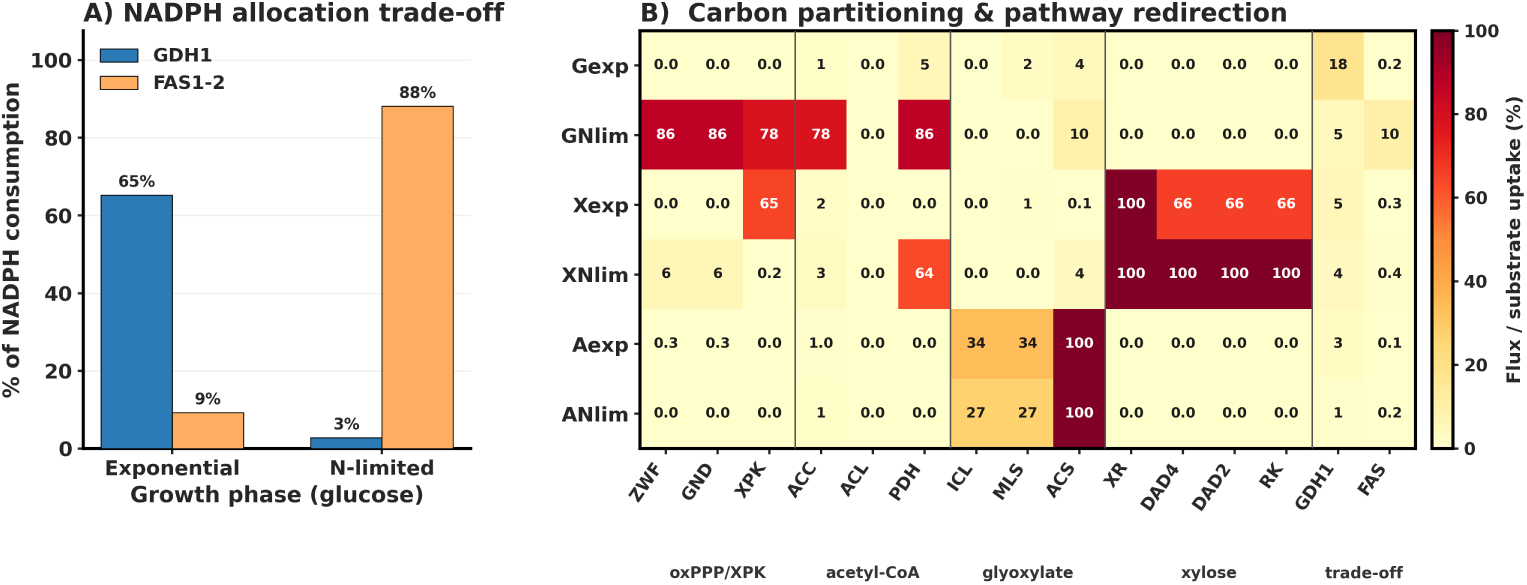
kinGEMs recovers major resource-allocation trade-offs in *Rhodotorula toruloides* across carbon sources and growth phases. **(A)** Percentage of cytosolic NADPH consumption attributed to NADP-dependent glutamate dehydrogenase (GDH1, blue) versus fatty acid synthase (FAS1–2, orange) during exponential and nitrogen-limited growth on glucose. **(B)** Predicted flux of selected marker reactions, normalized to the measured substrate uptake rate (%), for all six carbon-source × growth-phase combinations (Gexp/GNlim, glucose; Xexp/XNlim, xylose; Aexp/ANlim, acetate). Columns are grouped as oxidative pentose phosphate pathway and phosphoketolase (ZWF, GND, XPK); acetyl-CoA routes (ACC, ACL, PDH); glyoxylate / acetate entry (ICL, MLS, ACS); the D-arabinitol/D-ribulose xylose bypass (XR, DAD4, DAD2, RK); and the NADPH trade-off pair (GDH1, FAS). Each condition was annealed independently to its experimental growth-rate target under that condition’s medium, biomass composition, and enzyme-pool bound; intracellular path-way fluxes were not part of the objective (Methods 5.6.4).

On xylose, the clean GDH1 to FAS switch was not resolved (Fig 6B): both enzymes remained minor NADPH sinks in either phase (GDH1 ∼4–5%, FAS *<*1% of substrate-normalized flux). Instead, the NADP-dependent xylose reductase (XR) carried the entire xylose uptake in both phases and dominated NADPH demand. This is consistent with Rekena et al., who likewise found XR to be the single largest NADPH consumer on xylose, and reflects the stoichiometric NADPH cost of reducing xylose to xylitol at the entry point of the pathway. We report the unresolved GDH1–FAS trade-off on xylose as a genuine limitation, where the dominant XR sink masked the lipogenesis partition that is cleanly recovered on glucose.

#### 3.3.2 Carbon-source-specific pathway routing

Predicted fluxes recovered the substrate-specific gateways reported for *R. toruloides* (Fig 6B). Under glucose nitrogen limitation, oxidative pentose phosphate pathway flux (ZWF, GND; 86% of uptake) was coupled to phosphoketolase (XPK; 78%) and acetyl-CoA carboxylase (ACC; 78%), while ATP-citrate lyase (ACL) remained unused; this matches the acetyl-CoA route dependent on phosphoketolase that Rekena et al. identified in place of the PDH–CIT–ACL pathway. The one glucose discrepancy is the exponential-phase oxPPP, reported by Rekena et al. as constitutive at ∼72% of glucose carbon but not recovered here (ZWF/GND near zero in Gexp); in kinGEMs the oxPPP is engaged only under nitrogen limitation, when NADPH demand for lipogenesis peaks.

On xylose, flux used the alternative D-arabinitol/D-ribulose bypass (XR, DAD4, DAD2, RK) rather than xylulokinase, matching the absence of this enzyme in the experimental proteome. On acetate, acetyl-CoA synthetase carried the entire uptake and the glyoxylate shunt (ICL, MLS) carried 34% and 27% of uptake in exponential and nitrogen-limited phases, with PDH and ACL unused, reproducing acetate entry through ACS and the glyoxylate cycle. The one acetate route not recovered is cytosolic malic enzyme, reported by Rekena et al. as the sole NADPH source under acetate nitrogen limitation; its flux was negligible in all six conditions and it is therefore omitted from Panel B.

Taken together, from a single global enzyme-mass pool and *k*_cat_ values tuned only to growth rate, kinGEMs recovered the major qualitative allocation phenotypes of the manually curated, proteome-constrained reference. Predicted magnitudes differ from the Rekena et al. (sampled GECKO medians), and two reported features (constitutive exponential-phase oxPPP on glucose and cytosolic malic enzyme as the acetate nitrogen-limited NADPH source) were not reproduced. This provides direct evidence that kinGEMs is able to capture qualitative phenotypes in non-model organisms.

### 3.4 Emergent Recovery of Acetate Overflow Metabolism

Standard, non-enzyme-constrained FBA cannot predict acetate overflow (the shift toward partially fermentative metabolism observed in aerobic *E. coli* above a critical growth rate) without an artificial, externally imposed bound on oxygen uptake, since pure respiration is always the ATP-optimal strategy under unconstrained flux balance analysis at any growth rate. Enzyme-constrained models, by contrast, are expected to recover this behavior endogenously: as growth rate rises, a finite enzyme-mass budget forces a trade-off between the higher ATP yield but greater enzyme cost of respiration and the lower yield but cheaper enzyme cost of fermentation [33].

To test whether kinGEMs reproduces this emergent phenotype and, unlike the proteomics analyses above, to test generalization beyond a single *E. coli* reconstruction, we compiled literature-consensus growth rate, carbon uptake, and acetate secretion data for *E. coli* MG1655 across a range of aerobic growth conditions [33]. For three baseline GEMs spanning successive *E. coli* reconstructions (iAF1260, iML1515, iJO1366), we tuned *k_cat_* values via simulated annealing to match the reported maximum growth rate on minimal glucose medium, then simulated a range of growth rates by sweeping carbon uptake and recording predicted acetate secretion at each point (Fig 7).

**Fig 7:**
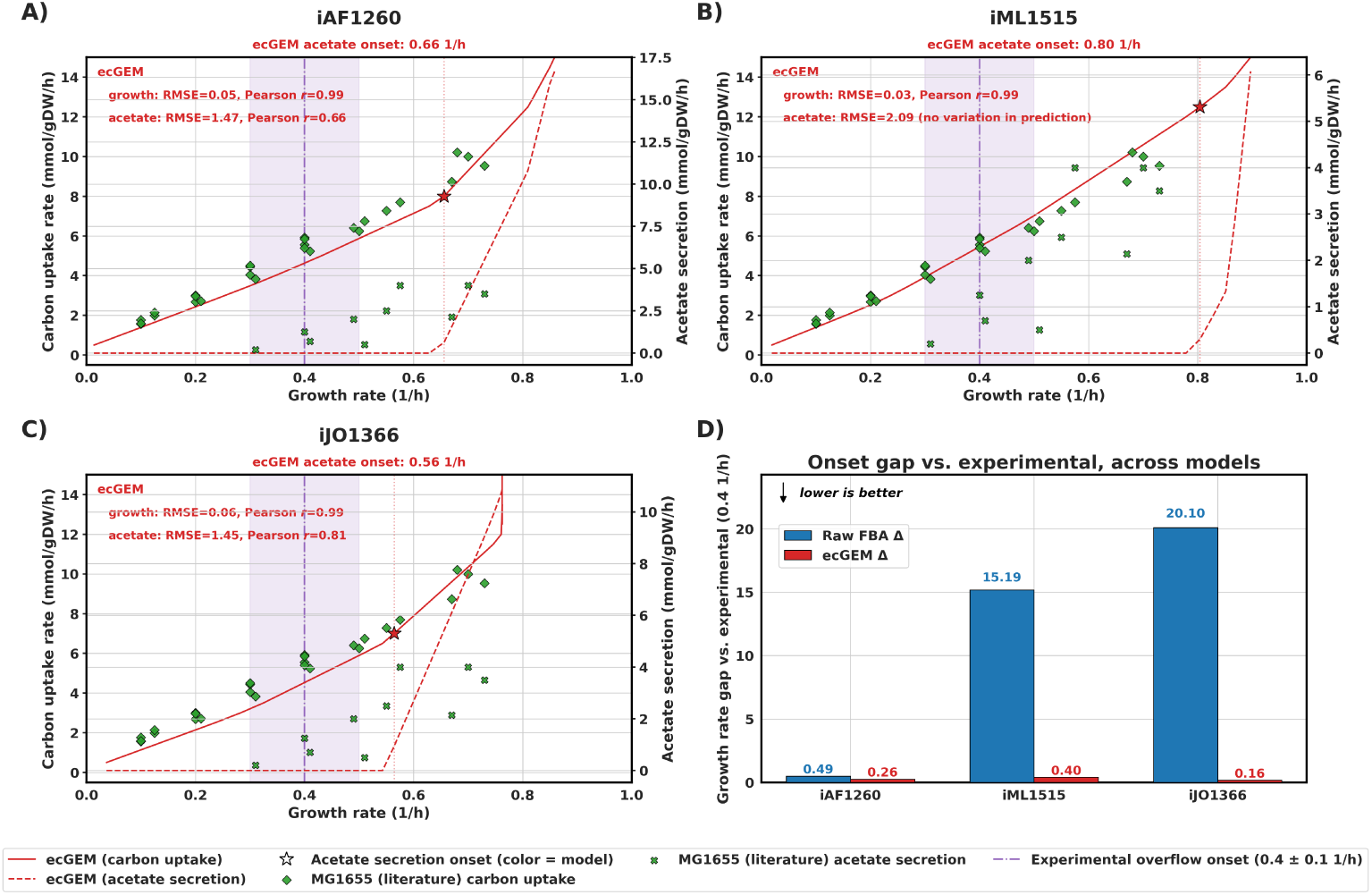
kinGEMs recovers emergent acetate overflow metabolism across three *E. coli* genome-scale reconstructions. **(A-C)** Carbon uptake rate (left axis, solid red) and predicted acetate secretion rate (right axis, dashed red) as a function of growth rate for enzyme-constrained (A) iAF1260, (B) iML1515, and (C) iJO1366 models, each tuned via simulated annealing to the experimentally reported maximum growth rate on minimal glucose medium. Green diamonds and crosses show literature-compiled *E. coli* MG1655 carbon uptake and acetate secretion data, respectively [33]. RMSE (mmol·gDW*^−^*^1^·h*^−^*^1^) is reported for both the carbon uptake fit (labeled “growth”) and the acetate secretion fit against these literature values; Pearson *r* is additionally reported where predicted acetate secretion varied sufficiently across the tested growth-rate range to support a correlation estimate (not defined for iML1515, whose predicted acetate secretion showed no appreciable variation over this range despite comparable RMSE to the other two models). Stars mark the model-predicted acetate secretion onset growth rate. The shaded band and dash-dot vertical line mark the experimental consensus overflow onset (0.4 ± 0.1 h*^−^*^1^); the dotted vertical line marks the ecGEM-predicted onset for each model. **(D)** Absolute gap between predicted and experimental acetate-onset growth rate (h*^−^*^1^) for raw, unconstrained FBA (blue) versus the enzyme-constrained kinGEMs model (red), compared across all three GEMs; lower is better. The enzyme-constrained models recover the experimental onset far more closely than the corresponding unconstrained baselines in every case (iAF1260: 0.26 vs. 0.49; iML1515: 0.40 vs. 15.19; iJO1366: 0.16 vs. 20.10).

All three enzyme-constrained models reproduced the qualitative relationship between carbon uptake and growth rate with high fidelity (iAF1260: RMSE = 0.05, *r* = 0.99; iML1515: RMSE = 0.03, *r* = 0.99; iJO1366: RMSE = 0.06, *r* = 0.99). More notably, each ecGEM predicted the onset of acetate secretion at a growth rate close to the experimentally observed consensus threshold of 0.4 ± 0.1 h*^−^*^1^ (iAF1260: 0.66 h*^−^*^1^, acetate RMSE = 1.47, *r* = 0.66; iML1515: 0.80 h*^−^*^1^, acetate RMSE = 2.09; iJO1366: 0.56 h*^−^*^1^, acetate RMSE = 1.45, *r* = 0.81), substantially outperforming the corresponding unconstrained FBA models, whose predicted onsets deviated from the experimental threshold by a much larger margin across all three GEMs (iAF1260: Δ = 0.49; iML1515: Δ = 15.19; iJO1366: Δ = 20.10, versus Δ = 0.26, 0.40, and 0.16 h*^−^*^1^ respectively for the enzyme-constrained models; Fig 7D). This indicates that the enzyme-mass budget introduced by kinGEMs is sufficient to recover a growth-rate-dependent metabolic switch that unconstrained stoichiometric models cannot capture without externally imposed bounds, providing phenotype-level evidence of biological realism beyond flux-interval precision or proteomic correlation alone.

We note one exception: unlike iAF1260 and iJO1366, the iML1515 ecGEM’s predicted acetate secretion showed no meaningful correlation with the literature-compiled data (Pearson *r* not reported, as predicted values did not vary sufficiently across the tested growth-rate range to support a correlation estimate), though its RMSE (2.09) remained comparable to the other two models (1.45-1.47), indicating the magnitude of error was similar even without a well-defined correlation. Growth-rate prediction for iML1515 remained strong (RMSE = 0.03, *r* = 0.99), and the model still recovered a qualitatively correct onset of acetate secretion close to the experimental threshold (0.80 h*^−^*^1^ vs. 0.4 ± 0.1 h*^−^*^1^ consensus). This indicates that while the onset behavior is robust across all three reconstructions tested, the graded shape of the secretion curve above onset is not uniformly well captured, and may depend on model-specific structural or parameterization differences that warrant further investigation.

## 4 Discussion

This work integrates machine learning-based kinetic prediction with automated construction and refinement of ecGEMs, providing a scalable route that improves both model *precision* and *accuracy*. By building models that demonstrate improvement along these two axes, we address a persistent model development and evaluation gap in the ecGEM space, where the emphasis is often on the prediction accuracy of enzyme kinetic parameters rather than the systematic assessment of downstream phenotype and flux-level consequences. Our results show that incorporating kinetic constraints estimated with CPI-Pred substantially reduces the feasible flux solution space, and that post hoc tuning of kinetic parameters can recover feasibility and improve agreement with experimental flux measurements while maintaining biologically realistic enzyme allocation constraints.

A central finding is that progressively integrating enzyme constraints yields reductions in flux variability, which clearly supports previous research that demonstrates that enzymatic constraints builds more precise GEMs by contracting the solution space [10, 21, 22, 34, 35]. However, tighter constraints also introduce a trade-off with achievable growth when kinetic parameters are imperfect or when model structure and proteome constraints are mismatched to the experimental condition. This trade-off highlights an important practical reality for ecGEM construction: **kinetic parameter prediction alone is insufficient if predictions induce systematically over-restrictive enzyme capacities for key pathways.** In this regime, the ability to *tune* parameters within relative uncertainty bounds is essential for reconciling mechanistic constraints with organism- and condition-specific phenotypes.

The simulated annealing framework implemented in kinGEMs provides a systematic approach to develop ecGEMs with biologically correct constraints. Rather than globally relaxing enzyme mass constraints via a single average saturation factor applied across the whole enzyme pool (as done prior approaches, e.g., GECKO’s sigma factor, which approximates the fraction of the total enzyme pool that is catalytically active [10, 21, 22]), kinGEMs targets the enzymes that dominate the global enzyme budget and perturbs their *k_cat_* values relative to the CPI-Pred-derived uncertainty ranges.

Consistent with this targeted, sparse-relaxation implementation and with strategies used by GECKO [21] and ECMpy [36], we find that restricting simulated annealing to a moderate perturbation pool (20-50 enzymes by mass contribution) achieves comparable or better feasibility and proteomics agreement at lower interpretability cost (Supplementary Information, Fig B7), suggesting that kinGEM’s tuning strategy could be made more sparse and interpretable without a meaningful loss of performance. This approach preserves a biologically plausible enzyme allocation while improving biomass feasibility and accuracy. Importantly, because the mapping from kinetic parameters to feasible growth is non-smooth and often includes infeasible regions, stochastic search offers a robust alternative to methods requiring differentiability or convexity assumptions. The result is an optimization loop that uses systems-level validation signals (growth in this study with the option of integrating other omics agreement setups) to refine molecular-level parameters while utilizing ML model-derived uncertainty.

Among constraint types, promiscuous enzyme formulations produced one of the largest reductions in flux variability. This supports the hypothesis that enzyme reuse across reactions is a major source of degeneracy in stoichiometric ecGEMs: when a shared enzyme pool must be allocated across competing pathways, many alternative flux modes are eliminated [37, 38]. This observation indicates that properly capturing enzyme sharing and competition is not just a biological nuance, but a fundamental structural factor governing model feasibility. It also motivates future work on improving annotation and inference of enzyme promiscuity, including systematic mapping of multi-substrate enzymes and isoenzyme specificity.

By taking a closer look at the enzymes perturbed during the simulated annealing process, we can identify which reactions and subsystems act as effective bottlenecks for achieving the target biomass (or other objective) in the ecGEM. Across all reactions, annealing produced a right-shifted, bimodal *k_cat_* distribution with a median increase from 5.6 to 54.3 (median fold-change ∼12.13×), with 51% of *k_cat_* values increasing and only 0.2% decreasing (Fig 8 A-D). This pattern indicates that, on average, the model compensates for under-estimated catalytic capacity rather than broadly down-scaling overly permissive enzymes, consistent with growth being limited by a relatively small subset of highly loaded reactions.

**Fig 8:**
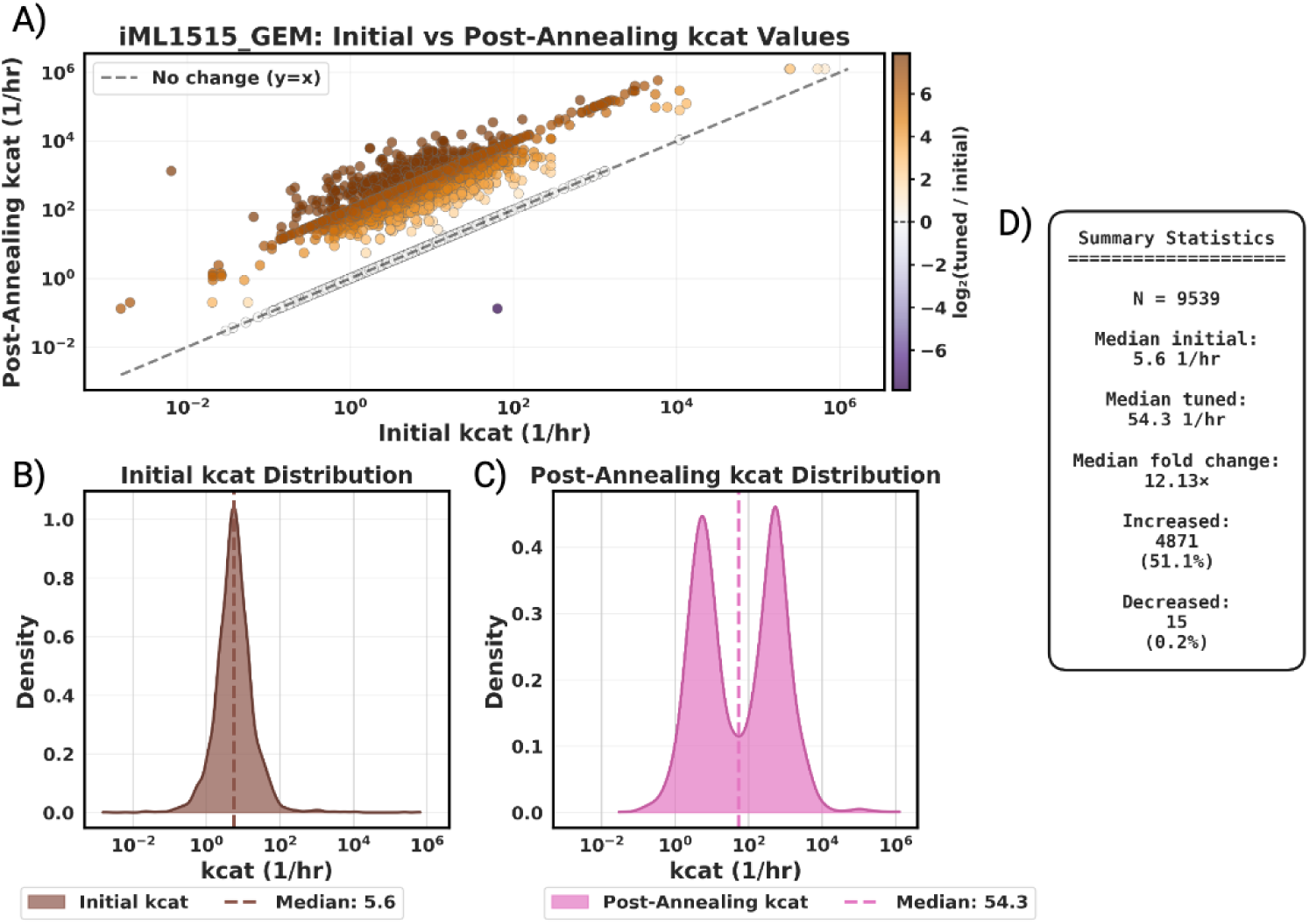
Global shifts in catalytic rate constants (*k_cat_*) following simulated annealing optimization. **(A)** Scatter plot comparing initial and post-annealing *k_cat_* values for all reactions in the iML1515 enzyme-constrained model. The dashed line indicates no change (*y* = *x*). Most points lie above the diagonal, indicating systematic increases in catalytic capacity. **(B)** Distribution of initial *k_cat_* values, showing a unimodal distribution centered at low turnover rates. **(C)** Distribution of post-annealing *k_cat_* values, exhibiting a rightward shift and increased variance, consistent with relaxation of overly restrictive enzyme constraints. **(D)** Summary statistics quantifying the global effect of annealing, including median *k_cat_*increase, fold-change, and proportions of reactions with increased or decreased catalytic rates.

Large fold-changes are concentrated in a targeted subset of enzymes rather than distributed broadly across the model. Perturbation is restricted to genes within the top 500 enzymes (per iteration) by mass contribution (of 1,516 total genes in iML1515); of these, 489 genes received an updated value. Nearly all of perturbed values increased, consistent with the algorithm’s stated bias towards positive perturbations (Section 5.4), and 37% of perturbed values reached the upper bound of the search space, a ceiling set at two orders of magnitude above the CPI-Pred-derived standard deviation. This indicates that a substantial share of the large fold-changes reported reflect enzymes saturating this explicit algorithmic bound rather than open-ended search escaping to biologically arbitrary values.

Decomposing these changes by subsystem reveals that the largest, most coherent upward shifts occur in membrane-associated and transport processes, including inner-membrane transport, outer-membrane porins, inorganic ion transport, and membrane lipid metabolism (Supplementary Fig B4). Subsystem-resolved comparison (Supplementary Fig B4) shows two distinct patterns: most subsystems (cofactor biosynthesis, alternate carbon metabolism, oxidative phosphorylation, nucleotide salvage, lipid and envelope biosynthesis) exhibit a clear bimodal structure, with a majority of reactions unperturbed and a smaller subset elevated 1–3 orders of magnitude, while outer-membrane porin transport (n=1,008) shows a broad, near-uniform upward shift across nearly the entire subsystem.This suggests that cross-membrane fluxes, which are already known to be strongly constrained by limited membrane area and protein crowding [33, 39], are potentially under-parameterized in the initial ML-based *k_cat_* assignment and therefore become primary targets for compensatory tuning.

However, systematic *k_cat_* increases reflecting underestimation of true catalytic capacity by the initial ML assignment is one plausible explanation among several that are not mutually exclusive. Because the simulated annealing procedure can compensate for multiple, distinct sources of model error through kinetic parameter adjustment alone, observed post-tuning *k_cat_* shifts should not be interpreted as direct evidence of true in vivo enzyme turnover rates. Alternative or contributing explanations include incomplete gene-protein-reaction mappings, missing regulatory constraints not captured by the stoichiometric formulation, and structural inaccuracies in enzyme allocation assumptions more broadly. We regard the membrane/transport pattern discussed above as a literature-supported and biologically plausible contributor to the observed shifts, rather than as the sole or confirmed explanation, and emphasize that the annealing procedure is fundamentally one of model-level compensation for whichever errors (kinetic, structural, or regulatory) prevent the model from matching the growth-rate objective.

One candidate mechanism specific to the membrane and transport pattern observed here related to enzyme reuse and promiscuity. From the perspective of enzyme reuse, these results imply that a small number of promiscuous or heavily shared transport proteins carry disproportionately large fluxes and thus dominate the global enzyme allocation budget. When a single transporter services multiple substrates or connects several systems, any underestimate of its *k_cat_* forces the annealing procedure to inflate its effective turnover to reconcile the shared capacity with the observed growth rate. In contrast, enzymes in more modular, pathway-specific segments of metabolism can often redistribute flux through alternative routes, so their *k_cat_* values require less drastic changes. Together, these patterns are compatible with the idea that accurately modeling enzyme sharing is critical for membrane and transport processes, where a few multi-use proteins couple many pathways and therefore have large control over both flux variability and feasible growth phenotypes, while the alternative sources of model error noted above remain equally viable explanations for the same observations.

Agreement between model-predicted feasible flux ranges and ^13^C MFA ranges improved after introducing enzymatic constraints, and improved further after tuning. Notably, the baseline GEM exhibits apparent agreement in some cases due to permissive solution spaces rather than mechanistic accuracy. In contrast, kinGEMs produces narrower predicted ranges that are more informative for hypothesis generation and strain design. At the same time, the sharp increase in zero-overlap reactions (Fig 4, bottom of panel B) after imposing constraints, followed by recovery after tuning, underscores a critical point: precision-enhancing constraints don’t necessarily lead to better alignment with experimental flux data, and therefore must be paired with tuning methods to adjust the parameters so as to eliminate systematic mismatches introduced by ML-predicted kinetics and model structure (missing or incorrect constraints, missing reaction-gene mapping, optimization solution alternates, etc.). These residual gaps present opportunities for future refinement and highlight the value of fluxomics as a rigorous validation layer beyond growth phenotypes.

While we demonstrate improvement in precision and accuracy on *E. coli* GEMs through this kinGEMs pipeline, the broader implication is that automated kinetic integration and tuning may produce greater accuracy improvements in less-curated organisms, where missing kinetics and enzyme allocation constraints represent a major source of uncertainty. kinGEMs is designed to operate exactly in this setting: it can construct ecGEMs from baseline models, populate missing kinetic parameters using CPI-Pred, and then reconcile predictions with organism-specific measurements through tunable refinement. As community-scale metabolic model resources continue to expand, this capability is important for enabling comparative systems biology and accelerating metabolic engineering in non-model organisms. To demonstrate the scalability of this pipeline, we ran the kinGEMs pipeline on the BiGG database [26] to provide *k_cat_* tuned and annotated ecGEMs for a total of 93 models (Supplementary Table E5). Since kinGEMs utilizes gene IDs stored in the GEMs and converts them to protein sequences as a data processing step prior to CPI-Pred’s inference stage, models that contain fragmented, deprecated, or incorrect gene IDs and could not be mapped to reference gene or protein sequences were not processed. Future work that addresses this gap of mapping GEM gene ID annotations to their respective proteins would allow for further expansion of the kinGEMs pipeline. For example, we could generate hundreds of thousands of kinGEMs models with efficient compute times (Supplementary Fig E12) on databases with thousands of models such as AGORA2 (7,302 models) [40] and APOLLO (247,092 models) [41].

Full statistics on the processed models can be found in the Supplementary Information Table E6. The models are available in the GitHub repository: https://github.com/LMSE/kinGEMs.

To assess the quality of these 93 models beyond aggregate feasibility, we examined how enzyme-constrained biomass recovery (ΔBM_EC vs FBA_, Supplementary Table E6) varies with phylogenetic distance from *E. coli*, and, for the subset of organisms with an available reference, how tuned growth rates compare against literature-reported maximum growth rates (Fig 9). *E. coli* /*Shigella* models (*n* = 64) showed the tightest and smallest median deviation from their unconstrained baseline (∼8%), and this deviation increased progressively with phylogenetic distance, reaching its largest median and widest spread among the 14 eukaryotic models in the set (Fig 9A). This pattern is consistent with CPI-Pred’s training data and the annealing procedure’s tuning behavior being most effective for the bacterial, and particularly enterobacterial, metabolism it was developed and validated on, and indicates that model quality for the 93-model set is not uniform across taxa, a limitation we discuss further below. A small number of individual models showed substantially larger deviations than their taxonomic group’s median (e.g., *S. aureus*, −99.3%; several *Trypanosoma cruzi* strain variants, −57 to −75%), which we attribute to a combination of sparser CPI-Pred sequence/kinetic coverage and, in several cases, substantially smaller model size (fewer genes and reactions) than the median model in this study.

**Fig 9:**
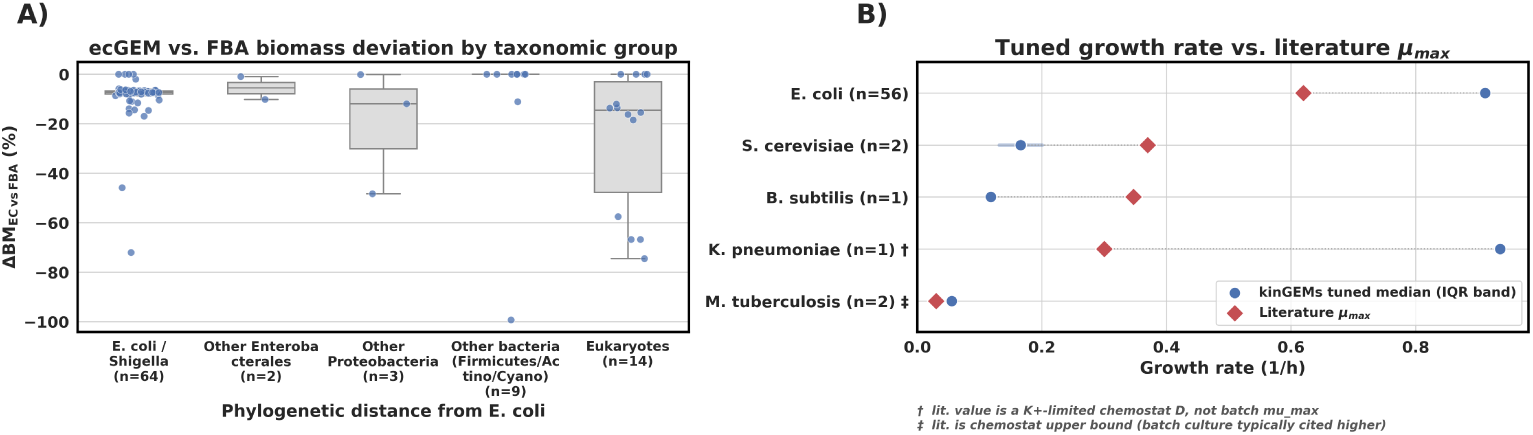
Quality of the 93 tuned BiGG ecGEMs by phylogenetic distance from *E. coli* and against literature growth rates. **(A)** Percent deviation of enzyme-constrained (tuned) biomass flux from the corresponding unconstrained FBA optimum (ΔBM_EC vs FBA_; Supplementary Table E6), grouped by phylogenetic distance from *E. coli* : *E. coli* /*Shigella* (*n* = 64), other Enterobacterales (*n* = 2), other Proteobacteria (*n* = 3), other bacteria (Firmicutes/Actinobacteria/Cyanobacteria, *n* = 9), and eukaryotes (*n* = 14). Boxes show interquartile range and median; points show individual models. Deviation from the unconstrained baseline increases with phylogenetic distance, with the eukaryote group showing both the largest median deviation and widest spread. **(B)** Simulated-annealing-tuned growth rate (blue) compared against literature-reported maximum growth rate (red diamond) for the subset of organisms in the 93-model set with an available literature reference: *E. coli* K-12 MG1655 [42], *S. cerevisiae* [43], *B. subtilis* [44], *K. pneumoniae* [45], and *M. tuberculosis* [46] (*n* per organism as labeled; points and bars show median and interquartile range across models for that organism). For *K. pneumoniae* (†), the literature value is a potassium-limited chemostat dilution rate (*D* = 0.3 h*^−^*^1^) rather than batch *µ*_max_; for *M. tuberculosis* (‡), the literature value is a chemostat-controlled growth rate rather than batch *µ*_max_. Both should be interpreted with this caveat rather than as direct evidence of over- or under-prediction.

Comparing tuned growth rates against literature-reported values for the five organisms in the set with an available reference (Fig 9B), *E. coli* models showed reasonable agreement with literature *µ*_max_ [42], while *S. cerevisiae* [43] and *B. subtilis* [44] models underpredicted growth rate relative to literature. For *K. pneumoniae*, the only available literature reference is a potassium-limited chemostat dilution rate (*D* = 0.3 h*^−^*^1^) rather than batch *µ*_max_ [45]; for *M. tuberculosis*, the available reference is likewise a chemostat-controlled growth rate rather than batch *µ*_max_ [46]. Because chemostat dilution rates are, by design, held below the organism’s true maximum growth rate, the apparent over-prediction for *K. pneumoniae* and close agreement for *M. tuberculosis* should not be read as direct evidence of model accuracy or inaccuracy for these two organisms. Taken together with the taxonomic pattern in panel A, these results indicate that the current kinGEMs pipeline, applied without organism-specific adjustment, produces its most reliable models for *E. coli* and closely related Enterobacterales, with quality degrading, though remaining broadly feasible, for more phylogenetically distant and eukaryotic organisms, underscoring both the value of the scalability demonstrated here and the importance of taxon-aware validation, such as the *Rhodotorula toruloides* analysis presented in Section 3.3, when extending this framework to non-model organisms.

In this study, we make several assumptions that are important to note when interpreting results. First, we primarily focus on integrating predicted *k_cat_* values, whereas other kinetic parameters (*K_M_* , *K_I_* , and *k_cat_/K_M_* ) can materially influence effective enzyme capacity *in vivo* through substrate availability and inhibition. Pipelines like COBRA-k [34] demonstrate the benefit of adding these additional enzymatic constraints within the modeling framework. Although CPI-Pred supports these parameters, incorporating them into genome-scale models may require nonlinear or approximated constraint formulations and condition-specific metabolite data, which can be incorporated in future iterations of this work. While our approach remains within the steady-state, linear-optimization paradigm, it is complementary to large-scale kinetic frameworks that leverage deep learning to reconstruct dynamic models from data-constrained parameter ensembles [47]. Second, enzyme constraints in ecGEMs typically approximate effective catalytic capacity using *in vitro*-derived or predicted parameters, which may not capture allosteric regulation, post-translational modification, compartmentalization, or condition-dependent enzyme usage [24]. Third, the tuning framework optimizes parameters to match selected system-level objectives (e.g., growth feasibility, flux consistency), which risks compensating for structural model errors by adjusting kinetics. This risk can be mitigated by multi-objective tuning against orthogonal datasets (proteomics, fluxomics, knockout fitness) and by restricting parameter adjustments for well-characterized enzymes.

The kinGEMs framework suggests several avenues for further validation and improvement. Experimentally, enzymes identified as dominant contributors to the global enzyme budget or those repeatedly adjusted during tuning provide a short list of high-impact targets for kinetic characterization. Targeted assays (e.g., purified enzyme turnover measurements under relevant substrate/cofactor conditions) or high-throughput approaches such as multiplexed enzyme activity screens could directly test whether tuned parameters move closer to condition-specific effective kinetics. At the systems level, additional ^13^C MFA datasets across different carbon sources and growth rates would enable more stringent evaluation of internal flux predictions and the transferability of tuned parameters across conditions. Computationally, integrating proteomics could constrain enzyme abundances directly, reducing reliance on global enzyme pool bounds and improving GEM-ecGEM differentiability. More broadly, incorporating metabolomics-informed saturation estimates or inhibition effects could bridge the gap between static kinetic parameters and context-dependent enzymatic capacity. Together, these directions would move ecGEMs toward mechanistically faithful, condition-aware models capable of guiding predictive metabolic engineering.

## 5 Methods

### 5.1 kinGEMs Pipeline

#### 5.2 Constraint Formulation for Enzyme-Constrained Models

The kinGEMs framework builds upon established ecGEM methodologies that incorporate enzyme kinetics and proteomics constraints [10, 21, 48]. GEMs are represented as systems of linear equations that describe metabolic fluxes, where stoichiometric constraints enforce mass balance. In ecGEMs, enzyme capacity constraints further restrict reaction fluxes based on enzyme abundances and catalytic efficiencies. The kinGEMs framework begins by parsing a baseline GEM to identify substrates and enzymes through gene-protein-reaction (GPR) associations. Subsequently, *k_cat_* values are predicted using CPI-Pred to address the sparse availability of experimental kinetic data, and these predictions are integrated into ecGEM to establish flux constraints.

The constraint formulation accounts for several biological complexities through mathematically rigorous representations (Table 2):

- **Simple scenario:** For reactions catalyzed by a single enzyme, the flux is constrained by the product of the enzyme’s turnover number and its concentration
- **Isoenzymes (OR logic):** When multiple enzymes can independently catalyze the same reaction, the total flux is constrained by the sum of individual enzyme capacities. This represents functional redundancy where any enzyme can contribute to the reaction at different points in time or simultaneously
- **Enzyme complexes (AND logic):** For reactions requiring multi-subunit enzyme complexes, the flux is limited by the least abundant subunit (the rate-limiting component). This constraint is expressed using a minimum function to capture the stoichiometric requirement
- **Promiscuous enzymes:** When a single enzyme catalyzes multiple reactions, the total enzymatic demand across all reactions cannot exceed the available enzyme pool These constraints are implemented using disjunctive normal form to handle complex GPR associations (detailed in the Supplementary Information Algorithm 1). Our disjunctive normal form treatment of GPR rules complements earlier formulations developed for efficient computation of gene-level minimal cut sets and intervention strategies [49, 50]. Additionally, a global constraint limits the total enzyme mass to biologically realistic values:

**Table 2:** Constraint formulations for different enzyme-reaction scenarios in kinGEMs.

| Scenario | Enzymes | Reaction | Flux Formulation |
| --- | --- | --- | --- |
| Baseline | single | single | $v_i \leq k_{cat,i} \cdot [E_i]$ |
| Isoenzymes (OR) | multiple | single | $v_i \leq \sum_{j=0}^J k_{cat,i,j} \cdot [E_{ij}]$ |
| Enzyme complex (AND) | multiple | single | $v_i \leq \min(k_{cat,i} \cdot [E_{ij}])$ |
| Promiscuous Enzymes | single | multiple | $\sum_{i=0}^I \frac{v_i}{k_{cat,i,j}} \leq [E_j]$ |

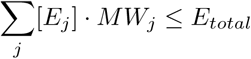

where *MW_j_* represents the molecular weight of enzyme *j*, and *E_total_* is the maximum allowable enzyme mass fraction of cellular dry weight. Our linear capacity constraints, adapted from earlier work [10, 51], can be viewed as a coarse abstraction of more detailed modular rate-law and convenience-kinetics formulations, which provide thermodynamically consistent expressions for multi-substrate reactions and regulatory effects [52]. All enzyme capacity constraints are formulated to preserve linearity, ensuring that the resulting optimization remains a linear program compatible with standard linear solvers. The pipeline, however, is built to allow for the integration of non-linear constraints, such as saturation and regulatory enzymatic constraints.

### 5.3 Integration of Kinetic Parameters and their Uncertainty

CPI-Pred predictions are obtained from 5-fold cross-validated ensemble models, yielding five independent *k_cat_* estimates for each enzyme-substrate pair. kinGEMs averages these predictions to obtain point estimates and calculates their standard deviations, which define *k_cat_* ranges for parameter uncertainty. These standard deviations subsequently inform the simulated annealing optimization [53] (described below), enabling exploration of the kinetic parameter space while maintaining consistency with the learned distributions from CPI-Pred.

### 5.4 Simulated Annealing for Kinetic Parameter Optimization

Once kinetic constraints are established, the model is solved using FBA with the objective of maximizing biomass production, though alternative objective functions can be specified for other applications. However, initial implementations revealed that enforcing total enzyme concentrations within experimentally observed ranges resulted in severely underestimated growth rates. Previous approaches [21, 22] address this by relaxing the total enzyme constraint to 50% of dry cell weight, corresponding to total protein rather than active enzyme content. Although this relaxation improves growth predictions, it makes the biologically unrealistic assumption that all cellular proteins are catalytically active, when realistically only about 10-25% of cellular mass comprises functional metabolic enzymes, with the total protein accounting for approximately 55% [54]. COBRA-k allows for the 25% total protein constraint, but does not consider genome-scale, automatic implementation and objective-directed tuning of these kinetic parameter values and is focused only on the *Escherichia coli* model [34].

To maintain biologically realistic enzyme allocations while improving predictive accuracy, kinGEMs implements a simulated annealing framework that iteratively refines *k_cat_* values with respect to CPI-Pred-derived uncertainty ranges (Fig 10). Treating kinetic parameters as distributions in this step, rather than fixed scalars, aligns with frameworks that generate ensembles of thermodynamically consistent kinetic models to capture uncertainty and variability in enzyme kinetics [47, 55–57]. The algorithm starts by computing the objective function value through the kinGEMs optimization module (biomass in all presented results). The enzymes are sorted by largest enzyme mass and the top 500 enzymes are then selected for further processing. We randomly sample 10-25% of these top 500 enzymes to perturb per iteration. In each iteration, the perturbation process starts by defining absolute lower bounds and upper bounds for the *k_cat_* values per enzyme, where both values are determined by 2 orders of magnitude of the standard deviation values coming from the 5-fold cross-validated CPI-Pred models’ predictions. In practice, 37.0% of perturbed enzymes reach this upper bound, indicating that the largest reported fold-changes are frequently a direct consequence of this explicit clamp rather than unconstrained exploration by the annealing search. In each iteration, we bias positive *k_cat_* perturbations over negative *k_cat_* perturbations, meaning that there is a higher chance that the *k_cat_* value will increase rather than decrease in each iteration. This allows for relaxation of the flux space to allow for higher biomass values from the overall optimization solution. Once the *k_cat_* values are updated for all the enzymes selected in the iteration, the optimization problem is rerun to obtain the new objective function value. If the value of the objective function improves (increase in biomass), the *k_cat_* values are automatically accepted. If there is no change or the objective function value regresses, a temperature function decides whether these newly obtained *k_cat_* values are carried out in future iterations or discarded. This algorithm allows for exploration earlier in the iteration processes and then exploits positive changes as it moves further in the process. The algorithm can be found in the Supplementary Information section B under Algorithm 2, and is also summarized in Fig 10. For the results presented in the main text, simulated annealing was run with the following configuration: initial temperature *T*_0_ = 100, geometric cooling rate *α* = 0.95 applied per iteration (*T* ← *αT* ), minimum temperature *T*_min_ = 0.001, maximum iterations = 1,000, and early stopping after 50 iterations without an improvement exceeding a 0.1% change threshold. Perturbation was restricted to the top 500 enzymes by mass contribution at each iteration, consistent with the enzyme-mass budget of 25% of dry cell weight described above.

**Fig 10:**
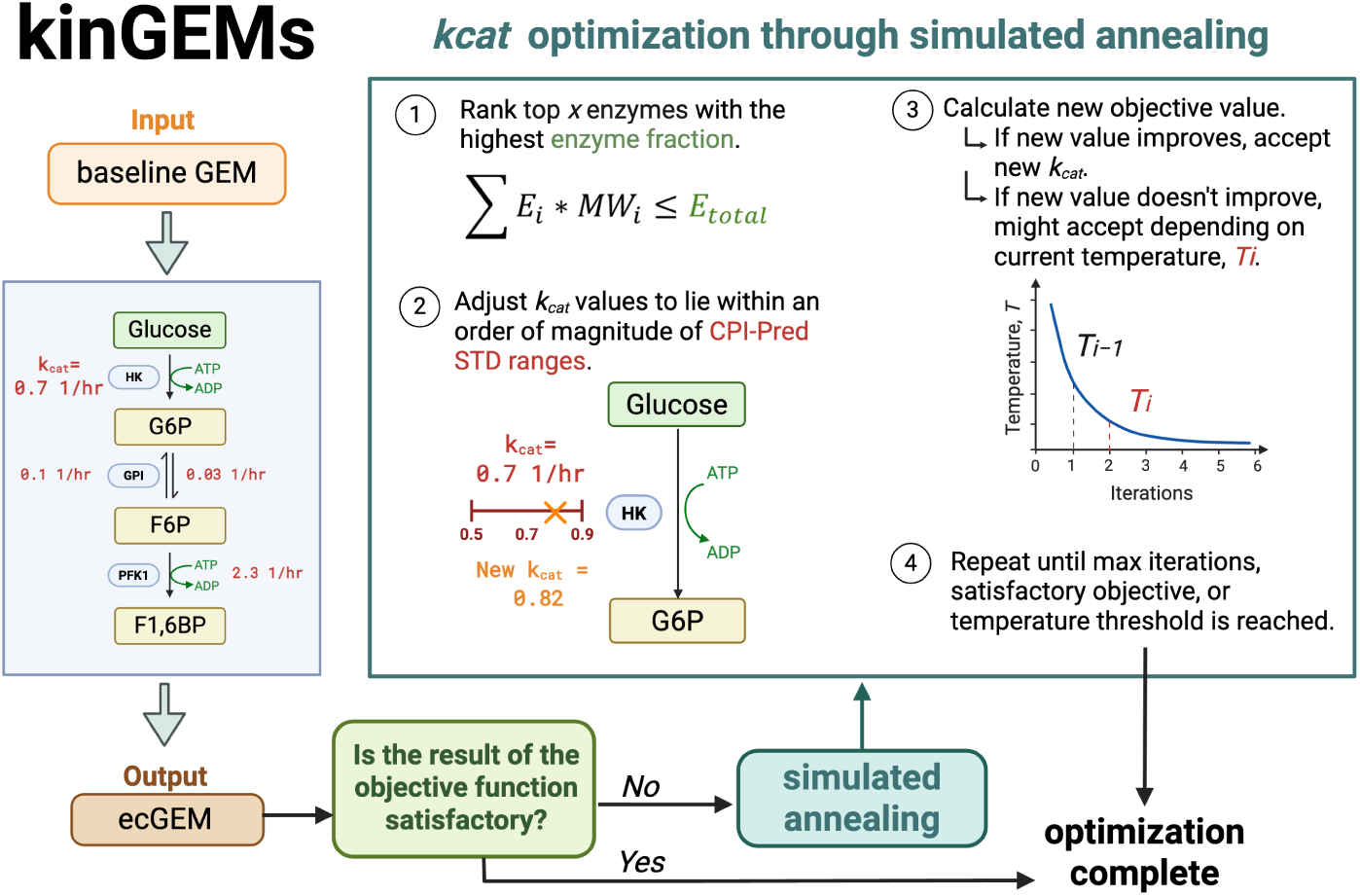
Simulated annealing framework for kinetic parameter optimization in kinGEMs. The optimization workflow begins with a baseline GEM as input and iteratively refines *k_cat_* values to improve model predictions while maintaining biologically realistic enzyme allocations. **Step 1:** Enzymes are ranked by their contribution to total enzyme mass ( *E_i_* · *MW_i_* ≤ *E_total_*), identifying the top *x* enzymes that most significantly constrain the model. **Step 2:** The *k_cat_* values of selected enzymes are randomly adjusted within relative CPI-Pred standard deviation ranges (e.g., *k_cat_* = 0.7 ± 0.2 1*/hr*), shown here perturbing from an initial value of 0.7 to a new value of 0.82 1*/hr*. **Step 3:** The objective function (typically biomass production) is recalculated. If the new value improves the objective, the updated *k_cat_* is accepted. If it worsens the objective, it may still be accepted probabilistically based on the current temperature *T_i_*, following the Metropolis criterion to escape local optima. **Step 4:** The temperature is gradually reduced according to a cooling schedule, and the process repeats until convergence criteria are met (maximum iterations, satisfactory objective value, or temperature threshold). The final output is an optimized ecGEM with improved predictive accuracy and realistic enzyme constraints.

Through this optimization algorithm, kinGEMs achieves biomass predictions that approximate the corresponding growth rates predicted by baseline (non-enzymatically constrained) GEMs. If further improvement is required, a maintenance parameter sweep over growth-associated maintenance (GAM) and non-growth-associated maintenance (NGAM) is performed to allow for growth rates that are consistent with experimental observations while maintaining realistic enzyme mass fractions (25% of dry cell weight). This sweep is a bounded sensitivity analysis rather than a free parameter search: for the iML1515 model, GAM was varied over {40, 45, 50, 55, 60, 70, 75.55, 80, 100} mmol ATP·gDW*^−^*^1^ and NGAM over {0, 1, 2*, . . . ,* 7} mmol ATP·gDW*^−^*^1^·h*^−^*^1^, a range bracketing the iML1515 literature-default values (GAM = 75.55 mmol ATP·gDW*^−^*^1^, explicitly included in the sweep grid; NGAM ≈ 3.15 mmol ATP·gDW*^−^*^1^·h*^−^*^1^ [27]), and its purpose is to characterize the sensitivity of predicted growth to maintenance-energy assumptions rather than to select a non-standard operating point; all biomass and flux results reported for iML1515 in this study use the literature-default GAM/NGAM values. The maintenance sweep was not performed for the remaining 92 BiGG models in this study, as *k_cat_*tuning alone was sufficient to recover growth consistent with baseline expectations; the same sweep methodology is available as an optional step in the kinGEMs pipeline for users seeking additional growth-rate recovery or wishing to adapt a model’s maintenance-energy parameterization to alternative experimental conditions. Importantly, this framework is extensible to incorporate additional experimental constraints, including fluxomics data, proteomic measurements of enzyme concentrations, or metabolomic profiles, which we demonstrate in Sections 3.3 and 3.2 enabling multi-omics integration for model refinement.

### 5.5 Flux Variability Analysis

To assess the impact of enzyme constraints on the solution space and evaluate model precision, we performed an FVA [58, 59]. FVA identifies the minimum and maximum flux values for each reaction while maintaining the optimal objective value (typically biomass production ≥ 90% of maximum). For each reaction *i*, FVA solves two optimization problems:

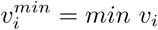

subject to:

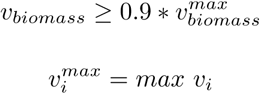

subject to:

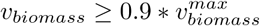

The flux variability for each reaction is then quantified as:

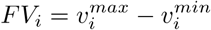

We compared flux variability distributions between baseline GEMs (stoichiometry-only constraints) and kinGEMs (enzyme-constrained) to quantify the reduction in solution space. Lower flux variability indicates tighter constraints and more precise flux predictions, which is critical for generating actionable hypotheses in metabolic engineering applications. We report median flux variability across all reactions and distributions.

To quantify how enzyme constraints introduced by the kinGEMs pipeline affect model precision, we assessed changes in the feasible flux solution space using FVA. We performed a systematic ablation study in which constraints were progressively added to a baseline GEM, followed by optional post hoc tuning of the kinetic parameters. For each constraint scenario, FVA was computed while enforcing near-optimal growth, with biomass flux constrained to the range

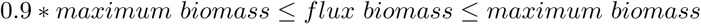

We evaluated seven levels of constraint complexity, summarized in Table 3: (L1) baseline GEM without enzyme constraints, (L2) single-enzyme constraints, (L3a-c) three independent single-constraint ablations relative to L2, each isolating one additional GPR-logic type ( isoenzymes, enzyme complexes, and promiscuous activity, respectively), (L4) all combined constraints, and (L5) all constraints with simulated annealing-based tuning of predicted *k_cat_* values.

**Table 3:** Progressive constraint hierarchy used in the FVA ablation study (Fig 3)

| Level | Constraints active | Description |
| --- | --- | --- |
| L1 | – | Baseline GEM; stoichiometric constraints only |
| L2 | Single-enzyme | Basic single-enzyme capacity constraints |
| L3a | + Isoenzymes | L2 + OR-logic isoenzyme constraints only |
| L3b | + Complexes | L2 + AND-logic enzyme-complex constraints only |
| L3c | + Promiscuous | L2 + promiscuous-enzyme coupling constraints only |
| L4 | All combined | All constraint types combined (complete kinGEMs) |
| L5 | All combined + SA | L4 model with simulated-annealing $k_{cat}$ tuning |

### 5.6 Benchmark Datasets

#### 5.6.1 Flux Variability Analysis for Fluxomics

To evaluate the alignment of internal flux predictions from kinGEMs against experimentally measured intracellular fluxes, we benchmarked against consolidated ^13^C MFA data, a gold-standard technique that integrates parallel ^13^C-labeling experiments to estimate central carbon fluxes with confidence intervals [60–62, 62–65]. We utilized a reference dataset that integrated 14 isotopic labeling experiments in *E. coli* into a consolidated metabolic network of 71 intracellular reactions and 22 exchange reactions [29]. Intracellular MFA reactions were manually assigned to their stoichiometric equivalents in the iML1515 GEM, resulting in 46 mappings (Table C1). To ensure a valid comparison, the GEM was constrained using the experimentally measured exchange fluxes reported in the MFA study. The lower and upper bounds of the exchange reactions in iML1515 were set to match the confidence intervals of the experimental data. The FVA was then performed as described in Section 5.5 to compute the feasible flux ranges.

Fixing exchange fluxes to Crown et al.’s experimentally measured values yields a lower FBA optimum under this constrained medium (0.4851 h*^−^*^1^) than the unconstrained-medium baseline used elsewhere in this study (0.87 h*^−^*^1^). Following simulated annealing (target: 0.75 h*^−^*^1^, the experimentally reported growth rate under these conditions [29]), a maintenance-parameter sweep was performed specifically for this fixed-medium analysis over the same GAM and NGAM ranges described above. The parameter combination minimizing deviation from the experimental growth target was NGAM = 10.0 mmol ATP·gDW*^−^*^1^·h*^−^*^1^ and GAM = 40.0 mmol ATP·gDW*^−^*^1^ (scaled down from the iML1515 default of 75.55), achieving a biomass of 0.504 h*^−^*^1^ (67% of the experimental target). This deviates from the literature-default GAM/NGAM values used in the general ablation and growth-recovery analyses in Fig 3; the adjustment reflects an empirical fit to this specific fixed-medium condition rather than a revised estimate of *E. coli* ’s general maintenance-energy requirements, and the residual gap to the experimental target likely reflects additional constraints (e.g., regulatory or condition-specific enzyme allocation effects) not captured by kinetic parameter and maintenance-energy tuning alone.

##### Precision of Flux Predictions

To evaluate the *precision* of internal flux predictions from kinGEMs against experimentally measured intracellular fluxes, we benchmarked kinGEMs model predictions against the consolidated ^13^C MFA data. In this analysis, precision was defined as the tightness of the feasible flux ranges under the imposed constraints. For each reaction *i*, we quantified the width of the FVA and MFA intervals as:

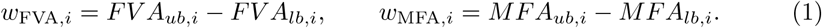

As summary measures of precision, we computed the average interval widths:

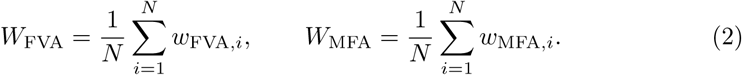

Smaller FVA interval widths (*W*_FVA_) indicate higher precision, as they reflect a narrower feasible flux range given the model constraints. Comparing *W*_FVA_ across unconstrained, GEM-constrained, and enzyme-constrained models quantifies how effectively additional mechanistic constraints increase precision by reducing solution space uncertainty. Comparison to *W*_MFA_ indicates how closely the model’s uncertainty matches experimental uncertainty.

##### Accuracy of Flux Predictions

In addition to precision, we assessed the *accuracy* of the predicted fluxes against the consolidated ^13^C MFA data. The agreement between the model predictions (FVA) and the experimental observations (MFA) was quantified using the *mean to mean distance*, which compares the central tendencies of the predicted and measured flux ranges.

For each reaction *i*, the midpoint of the FVA and MFA flux ranges is calculated as:

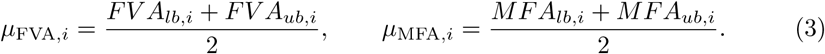

The absolute distance between these midpoints is then calculated as:

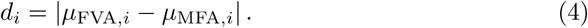

The overall mean-to-mean distance is defined as the average of these absolute differences across all evaluated reactions.

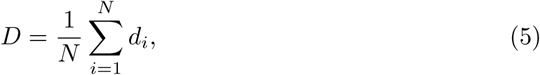

where *N* denotes the number of reactions for which both FVA and MFA bounds are available.

A lower value of *D* indicates stronger agreement between the model predictions and the experimental measurements, since it reflects a closer alignment between the central tendencies of the predicted and observed flux ranges.

The objective of this comparison is to determine whether the mechanistic models can produce a flux state consistent with experimental observations and to quantify how effectively enzymatic constraints narrow the feasible solution space compared to an unconstrained model.

#### 5.6.2 Proteomics Validation Against Absolute Quantitative Data

To evaluate agreement between predicted enzyme abundances and experimentall measured protein abundance, we benchmarked kinGEMs predictions against the absolute quantitative proteomics dataset of Schmidt et al. [30], spanning 15 *E. coli* growth conditions (glucose batch and chemostat cultures at four dilution rates; acetate, glycerol, pyruvate, fumarate, glucosamine, xylose, mannose, galactose, succinate, and fructose as sole carbon sources) and four independent *k_cat_* initialization sources (CPI-Pred; an independent deep learning predictor, CatPred; literature-derived BRENDA values; and randomly sampled log-normal values).

For each gene, predicted enzyme mass was computed from the completed enzyme-constrained FBA solution as enzyme mass = *v_E_* · *MW* × 10*^−^*^3^, where *v_E_* is the enzyme-usage flux variable and *MW* is the protein molecular weight (computed from sequence via Biopython). Because predicted enzyme mass (g protein/gDW biomass) and Schmidt et al.’s measured mass (fg/cell) do not share a common absolute scale, comparisons were performed in log10 space after fitting a single per-condition additive offset (the median of log_10_(predicted) − log_10_(measured) across matched genes), which was subtracted before computing RMSE and MAE; Pearson *r* is offset-invariant and was computed directly on log10 values without this correction, reflecting whether predicted and measured abundances rank genes consistently rather than whether they agree in absolute magnitude. Genes were matched between model gene identifiers (b-numbers) and Schmidt et al.’s UniProt accessions. When comparing across the four *k_cat_* sources for a given condition, RMSE and MAE were computed only on the intersection of genes with a valid, matched value across all four sources, ensuring a consistent gene basis for comparison; Pearson *r* was computed per source independently.

Because single-enzyme *k_cat_* prediction methods (CPI-Pred, BRENDA lookup, Cat-Pred) are not designed to characterize obligate multi-subunit enzyme complexes or translation machinery, we additionally verified proteomics agreement after excluding genes belonging to such categories, identified directly from iML1515’s own gene-protein-reaction rules (271 of 1516 genes, 17.9%): obligate subunits (GPR AND-clauses of size ≥ 2) of reactions in oxidative phosphorylation, transport (inner and outer membrane), and outer membrane porins (e.g. ATP synthase, NADH dehydrogenase, cytochrome oxidase), plus aminoacyl-tRNA synthetases (a near-no-op in this model, as iML1515 does not represent ribosomal proteins, RNA polymerase, or translation factors as metabolic genes).

#### 5.6.3 Cross-Condition and Pathway-Level Generalization of Tuned Kinetic Parameters

##### Cross-condition transfer

To test whether simulated-annealing-tuned *k_cat_* vectors generalize beyond the specific condition used to tune them, we performed a cross-condition transfer analysis. For each of the 15 growth conditions and 4 *k_cat_* sources, the growth-rate-tuned *k_cat_* vector (Section 5.4) produced under one (source) condition was applied, without further tuning, to every other (target) condition’s medium; FBA was re-solved under the target medium using this fixed *k_cat_* vector, and the resulting predicted enzyme abundances were scored against the target condition’s own measured proteomics using the procedure in Section 5.6.2. This yielded 767 cross-condition (source ̸= target) comparisons alongside 55 native (source = target) comparisons.

##### Proteomics-aware pathway-level tuning

As a complementary generalization check restricted to the glucose condition, we implemented a second simulated annealing phase that continues from the growth-rate-tuned (Phase 1) *k_cat_* vector but optimizes a different objective: the Pearson correlation (log10 scale) between predicted and measured proteomics, computed only over a training gene subset drawn from central energy metabolism (glycolysis, TCA cycle, pentose phosphate pathway, oxidative phosphorylation, and pyruvate metabolism). A disjoint, held-out test gene subset (ten amino acid biosynthesis pathways) was never included in the Phase 2 objective and was used only to evaluate whether proteomics-aware tuning generalizes to genes it did not directly optimize.

Phase 2’s perturbation rule differs from Phase 1’s: perturbations are symmetric (equal probability of increasing or decreasing *k_cat_*, as opposed to Phase 1’s bias toward increases), since improving proteomics fit requires moving some genes’ *k_cat_* values down as well as up. Each iteration perturbs 15-35% of train-pathway genes plus a small 5% dose of other eligible genes, rather than the full set simultaneously; an earlier formulation perturbing all eligible genes at once was found to rarely propose an improving move, as large simultaneous perturbations in a high-dimensional space are unlikely to improve the objective by chance and tend to break feasibility. A proposed move is accepted only if the resulting biomass remains within a tolerance (default 5%) of Phase 1’s own achieved biomass (not the original growth-rate target), since anchoring this tolerance to the growth-rate target rather than Phase 1’s actual outcome was found to reject nearly all proposed moves under enzyme budgets where Phase 1 substantially overshoots its target.

Statistical significance of the resulting change in Pearson *r* (Phase 2 minus Phase 1) was assessed for both the training and held-out test gene subsets using 95% bootstrap confidence intervals (percentile method, resampling matched gene pairs). Phase 2 improves fit on the training pathway by construction, since only proteomics-fit-improving moves are accepted; improvement on the held-out test pathway was not statistically significant in any completed run, indicating that proteomics-aware tuning under this procedure fits the specific pathway it is trained on without demonstrating reliable generalization to a distinct, unseen metabolic pathway within the same condition.

#### 5.6.4 Analysis of metabolic trade-offs in *Rhodotorula toruloides*

To test whether kinGEMs recovers intracellular resource allocation beyond growth rate, we reconstructed enzyme-constrained models of the oleaginous yeast *Rhodotorula toruloides* for the six carbon-source × growth-phase combinations reported by Rekena et al. [32] and compared predicted cofactor and carbon routing against their GECKO-based reconstructions. That study constrained every reaction with condition specific per-enzyme abundance bounds, BRENDA *k*_cat_ values, and manual *k*_cat_ curation. kinGEMs instead used a single global enzyme-mass pool, CPI-Pred kinetic predictions, and simulated annealing tuned to growth rate alone, with no proteomic or intracellular-flux data in its objective.

Six independent models were built using the rhto-GEM [31]. Each was constrained with the corresponding measured exchange rates. Condition specific protein, lipid, and carbohydrate mass fractions were imposed on the biomass pseudoreaction so that nitrogen-limited models were required to meet the experimental protein to lipid shift (values available in Table D3 and Table D4). Growth-associated ATP (GAM = 130 mmol ATP gDW*^−^*^1^) and non-growth-associated maintenance (NGAM = 3.65 mmol gDW*^−^*^1^ h*^−^*^1^) were applied before enzyme-constrained optimization and held fixed during annealing. Proteomics in Rekena et al. detected no xylulokinase. Following their reconstruction, the alternative fungal D-arabinitol/D-ribulose pathway was added, and xylulokinase was knocked out.

#### 5.6.5 Bootstrap annealing at experimental enzyme pools

The tight experimental enzyme pools (ranging from *E*_tot_ = 0.03 g gDW*^−^*^1^ to 0.09 g gDW*^−^*^1^) made the enzyme-constrained FBA infeasible. Simulated annealing cannot start from an infeasible point, because the enzymes to perturb are ranked by mass in an initial feasible solution. Whenever enzyme-constrained FBA at the experimental *E*_tot_ returned no feasible solution, we used a two-stage bootstrap:

1. *Seed solve.* Enzyme-constrained FBA was solved at a relaxed pool *E*_seed_ = 0.25 g gDW*^−^*^1^ to obtain a feasible solution.
2. *Stage 1 — restore feasibility at the experimental pool.* Simulated annealing was run at *E*_seed_ without stopping at *µ*_goal_, in chunks of 10 iterations. After each chunk, the tuned *k*_cat_ set was re-solved at the experimental *E*_tot_. Stage 1 ends at the first round for which the experimental pool becomes feasible.
3. *Stage 2 — match growth at the experimental pool.* Annealing was then continued at the experimental *E*_tot_ until *µ* ≥ *µ*_goal_ or the limits were reached.

#### 5.6.6 Emergent Recovery of Acetate Overflow Metabolism

To test whether kinGEMs recovers growth-rate-dependent acetate overflow without an explicit fitting objective for exchange fluxes, we compiled literature-consensus *E. coli* MG1655 growth rate, carbon uptake, and acetate secretion data across a range of aerobic growth conditions [33]. For three baseline GEMs spanning successive *E. coli* reconstructions (iAF1260, iML1515, iJO1366), enzyme constraints were applied and *k_cat_* values tuned via simulated annealing (Section 5.4) to match the reported maximum growth rate for each strain on minimal glucose medium. Using each resulting tuned model, we then simulated a range of growth rates by sweeping the glucose uptake rate and recording the corresponding predicted acetate secretion flux (EX ac e) at each point. Model fit to the literature carbon-uptake and acetate-secretion curves was assessed using RMSE (mmol·gDW*^−^*^1^·h*^−^*^1^) and, where predicted values showed sufficient variation to support a correlation estimate, Pearson *r*. The acetate-secretion onset growth rate for each model was defined as the lowest tested growth rate at which predicted acetate secretion first exceeded a small threshold above zero, and compared against the corresponding onset predicted by unconstrained FBA on the same baseline GEM, and against the experimental consensus onset of 0.4 ± 0.1 h*^−^*^1^.

#### 5.6.7 Sensitivity of Simulated Annealing to Perturbation Pool Size

To assess how the number of enzymes eligible for perturbation during simulated annealing (Section 5.4) affects feasibility, proteomics agreement, and interpretability, we repeated the growth-rate-tuning procedure with the eligible pool size *n_top_enzymes* ∈ {5, 10, 20, 50, 500} (CPI-Pred initialization, 8 growth conditions, 4 replicates per condition and pool size), spanning a sparse (GECKO/ECMpy-style) to broad (DLKcat-style, the pool size used for all other results in this study) regime. For each pool size, we recorded the percentage of the target growth rate reached, proteomics RMSE and Pearson *r* against Schmidt et al. proteomics (Section 5.6.2), and the number of genes whose *k_cat_* changed by more than ∼2.3% (a threshold distinguishing genuine perturbation from numerical noise) as a measure of interpretability cost.

## 6 Conclusion

We presented a unified framework that couples machine learning-based kinetic prediction (CPI-Pred) with automated construction and refinement of enzyme-constrained genome-scale models (kinGEMs). By benchmarking ecGEMs along two complementary axes; *precision*, measured by contraction of feasible flux ranges, and *accuracy*, measured by agreement with experimental flux and phenotype data, we provide a practical evaluation methodology for assessing the downstream impact of predicted kinetics in constraint-based modeling.

Across an ablation of constraint formulations, enzyme-informed constraints substantially reduced flux variability, demonstrating increased precision via a tighter and more informative solution space. Flux-level validation against ^13^C MFA further showed that enzymatic constraints, particularly when combined with parameter tuning, improve overlap with experimentally measured flux ranges. To address feasibility and biological realism, we introduced a simulated annealing approach that tunes predicted *k_cat_* values within relative CPI-Pred-derived uncertainty bounds while maintaining realistic limits on cellular enzyme allocation. This tunability enables ecGEMs to reconcile imperfect molecular predictions with organism- and condition-specific system-level constraints without relying on globally relaxed and biologically implausible enzyme mass assumptions.

Together, these results support the view that precision is a prerequisite for actionable genome-scale predictions, but that precision-enhancing constraints must be paired with methods for uncertainty-aware refinement to achieve accuracy. kinGEMs provides a scalable path for constructing and validating ecGEMs in both model and non-model organisms, and establishes a foundation for future integration of additional kinetic parameters and multi-omics constraints to build precise, condition-aware, mechanistic whole-cell models.

## 7 Supporting Information Legends

- Appendix A: kinGEMs Pipeline

– **Fig A1:** kinGEMs Process Overview
– **Algorithm 1:** Deriving enzyme-capacity reaction constraints from GPR rules (DNF-based)
– **Fig A2:** Flux variability ablation study
- Appendix B: Tuning ecGEMs Towards Experimental Alignment

– **Fig B3:** Optimization of biomass production during simulated annealing of kinetic parameters
– **Algorithm 2:** Simulated Annealing for Enzyme *k*_cat_ Optimization
– **Fig B4:** Subsystem-specific shifts in catalytic rate constants (*k*_cat_) following simulated annealing optimization
– **Fig B5:** Sensitivity of predicted growth rate to growth-associated (GAM) and non-growth-associated (NGAM) maintenance parameters in *E. coli* iML1515
– **Fig B6:** Reproducibility and convergence of simulated annealing optimization across 10 independent runs in *E. coli* iML1515
– **Fig B7:** Sparse vs. broad *k*_cat_ relaxation: effect of perturbation pool size on feasibility, proteomics fit, and interpretability
- Appendix C: MFA Experimental Comparison

– **Fig C8:** FVA Results Normalized Against Experimental MFA Values
– **Table C1:** Mapping between ^13^C MFA for intracellular and exchange reactions and iML1515 model reactions
– **Table C2:** Wilcoxon signed-rank tests comparing Baseline GEM against kinGEMs (post-tuning) and kinGEMs (pre-tuning) for mean-to-mean flux distance and FVA interval width (log-scale)
- Appendix D: Supplementary Proteomics and Phenotype Validation

– **Fig D9:** Predicted vs. experimental growth rate across 15 growth conditions and 4 *k*_cat_ sources
– **Fig D10:** Proteomics-aware tuning generalizes from trained to held-out metabolic pathways
– **Fig D11:** Cross-condition transfer of simulated-annealing-tuned *k*_cat_ vectors is comparable to native performance
– **Table D3:** Constraints for the *R. toruloides* reconstructions
– **Table D4:** Exchange bounds for the *R. toruloides* reconstructions
- Appendix E: Running kinGEMs on the BiGG Database

– **Table E5:** Aggregate statistics for the kinGEMs pipeline applied across 93 completed BiGG genome-scale metabolic models spanning 23 organisms
– **Fig E12:** Computational scalability of the kinGEMs pipeline across 93 BiGG genome-scale metabolic models spanning 21 organisms
– **Table E6:** Summary statistics across genome-scale metabolic models (*n* = 93 complete runs out of 94 attempted)

## Appendix A kinGEMs Pipeline

The kinGEMs pipeline is constructed to take in a pre-existing GEM as input along with a config file (in JSON format) with specifications for modeling parameters like total enzyme concentration as well as simulated annealing hyperparameters. The pipeline starts by reading the SBML file of the input GEM and extracting the SMILES representations of all the metabolites in the model through multiple SMILES-retrieval databases (Steps 1 and 2 in Fig A1). The pipeline also retrieves all the enzyme sequences in the GEM through the available gene IDs for all the reactions.

**Fig A1:**
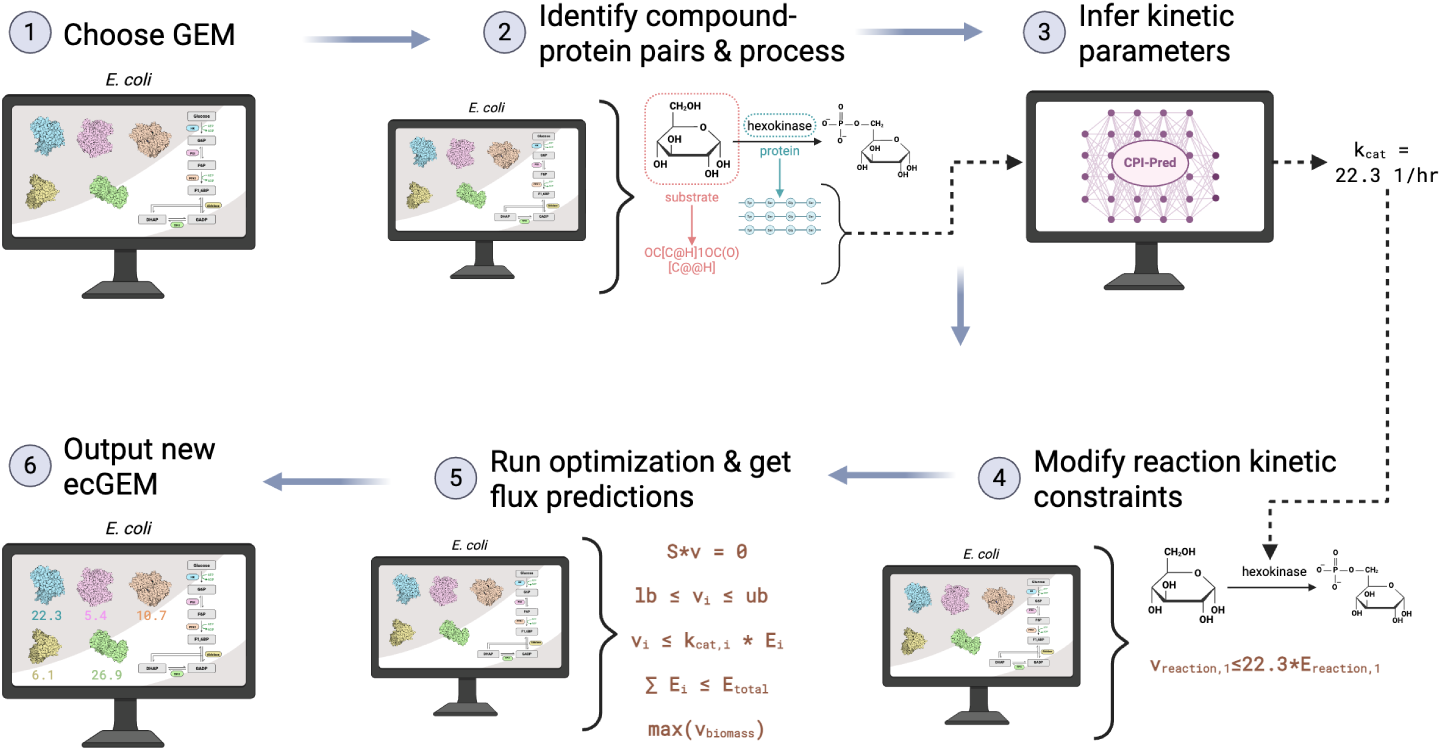
kinGEMs Process Overview. 1. The kinGEMs pipeline starts with a preexisting GEM as an input. **2** The compound-protein pairs in the entire GEM are then identified and processed. The SMILES representation of all the metabolites in the cell are obtained while the genes are used to identify and retrieve the corresponding protein sequence. **3** These compound-protein pair SMILES and sequences are sent to CPI-Pred where model inference is run to predict *k_cat_* values for all the pairs. **4** The *k_cat_* values are processed and mapped to specific reaction constraints through the GPR rules. **5** The optimization problem is run with all corresponding constraints and flux predictions are obtained. Simulated annealing is also run in this step. **6** The final annotated ecGEM is outputted and can be used for further downstream analyses.

The SMILES and protein sequence of every metabolite-enzyme pair is sent to the CPI-Pred model for inference. The model predicts *k_cat_* values for each pair with an associated standard deviation value (Step 3 in Fig A1). The standard deviation value is a product of obtaining five different predictions from five separate cross-validated CPI-Pred models. The pipeline then modifies the reaction constraints with the obtained *k_cat_* values through extracting and parsing the gene-protein-reaction rules in the GEM. This process is discussed in more detail in Algorithm 1.

Following the set up of the optimization problem with all the constraint conditions, the pipeline utilizes the Pyomo optimization framework [66] to run a linear programming problem with a maximization objective for the biomass (Step 5 in Fig A1). There are also intermediate optional steps pertaining to the simulated annealing process discussed in the Methods section. The pipeline then saves these newly built ecGEMs, which can be used in other types of analyses.

**Algorithm 1.** Deriving enzyme-capacity reaction constraints from GPR rules (DNF-based).

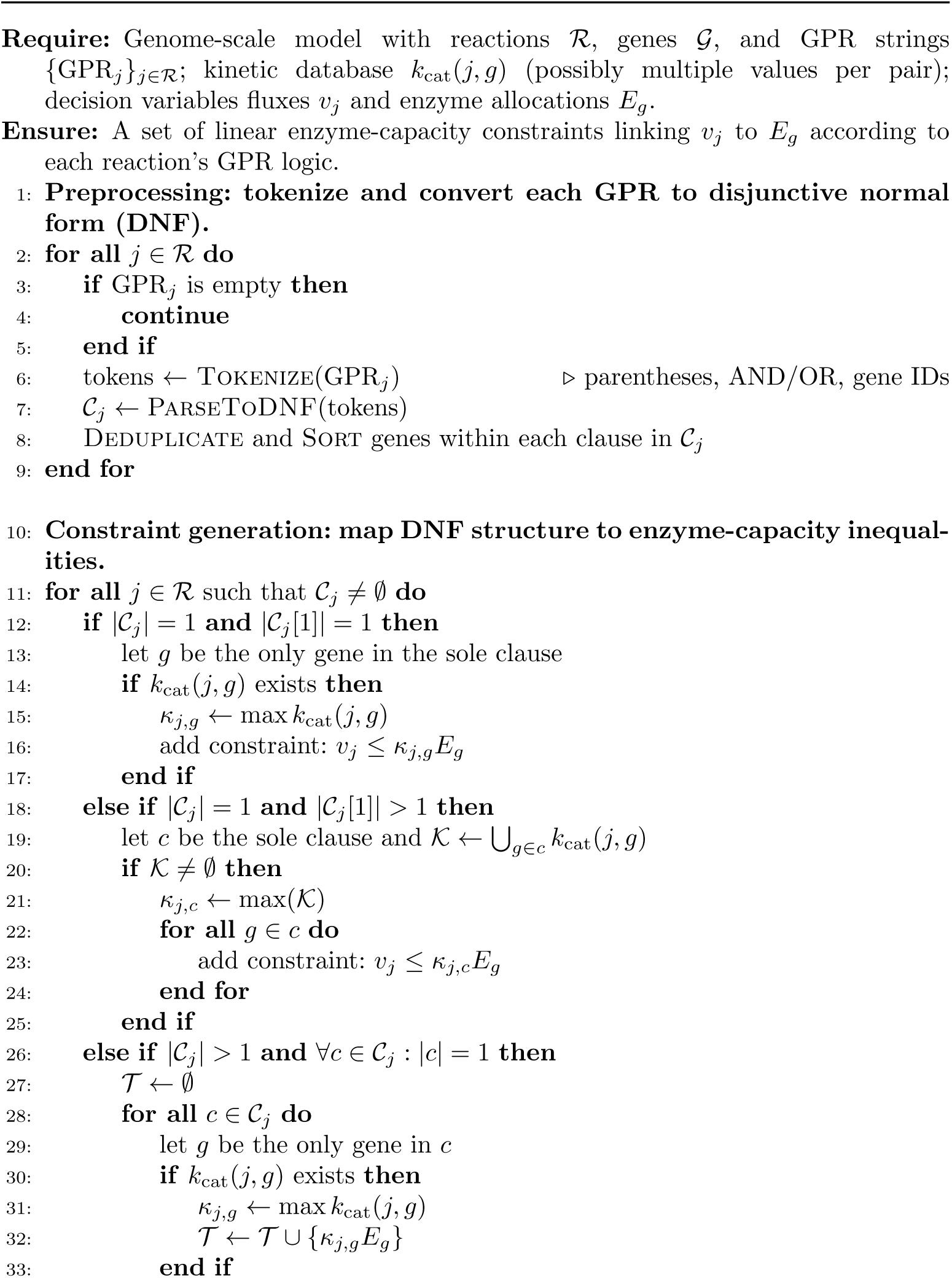
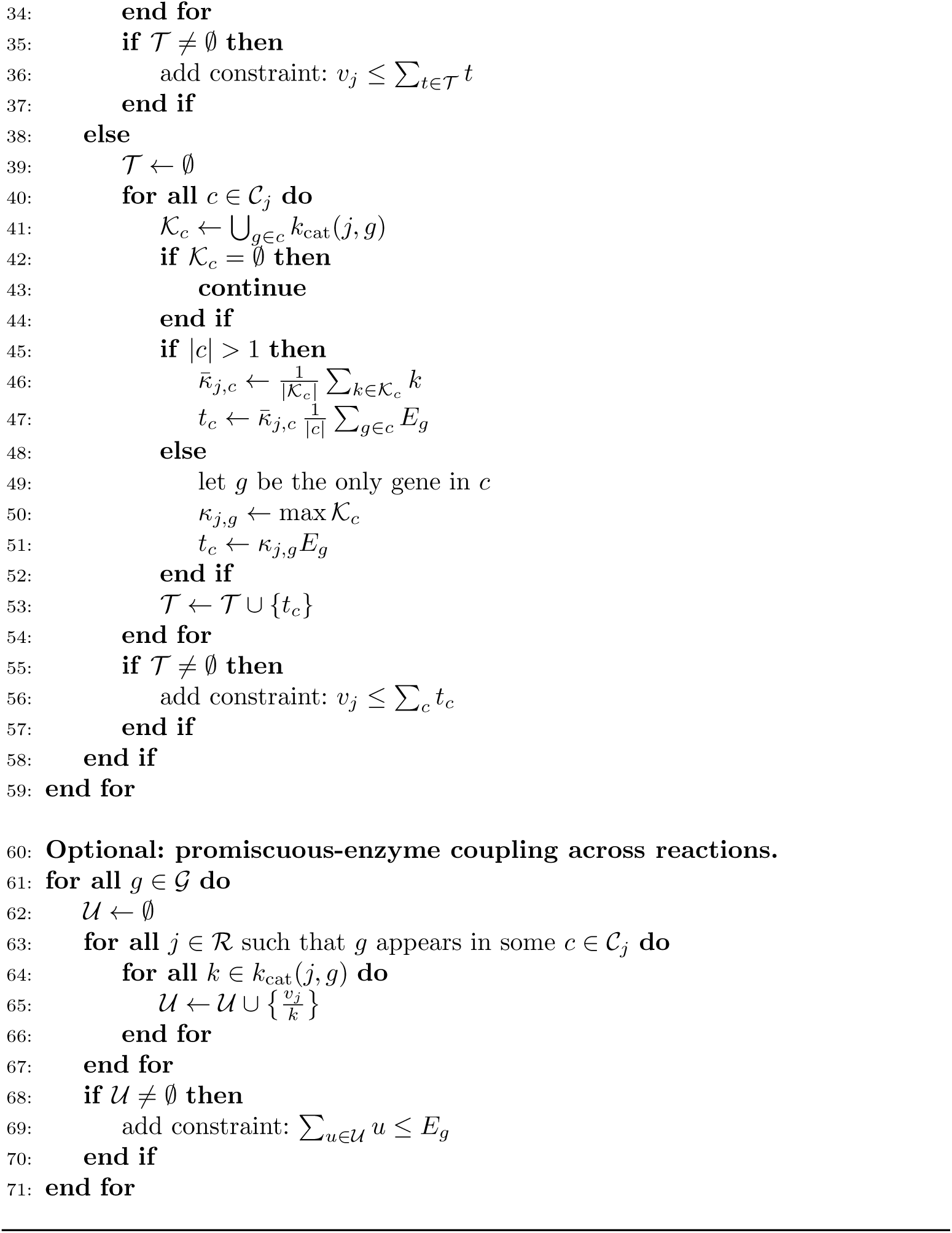

**Fig A2:**
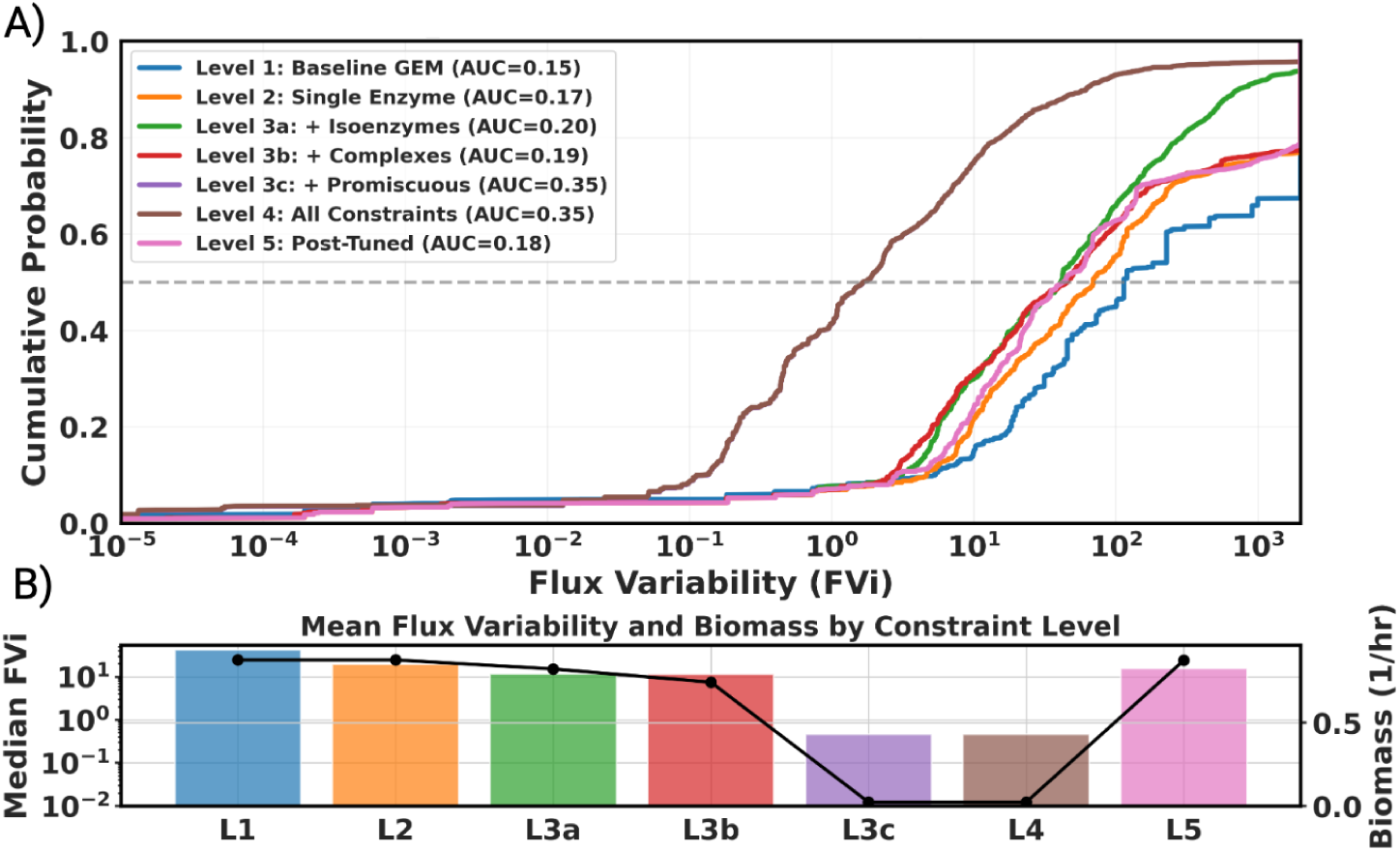
Flux variability ablation study. Progressive integration of enzyme constraints systematically reduces the feasible **non-optimal** flux solution space. **A)** Cumulative distribution probability of flux variability (FVi) across all reactions for each constraint level. Leftward shifts of the curves indicate tighter feasible flux ranges for an increasing fraction of reactions as enzymatic constraints are added. The dashed line denotes the median FVi. Here, FVi = *v*^max^ − *v*^min^ denotes the flux variability of reaction *i*. AUC here denotes area under the curve, where a higher value indicates greater proportion of reactions with low flux variability. *The level 3c line in the plot overlaps with Level 4, and is therefore not clearly visible in the plot. **B)** Median flux variability (bars, left axis) and corresponding optimal biomass flux (points, right axis) for each constraint level. While stronger constraints substantially reduce flux variability, they introduce trade-offs with achievable growth. Post hoc tuning of kinetic parameters (L5) restores biomass production while maintaining lower variability than L1.

## Appendix B Tuning ecGEMs Towards Experimental Alignment

As part of the kinGEMs pipeline, we present a simulated annealing algorithm that tunes the ML-predicted *k_cat_* values to better align them with experimental observations. In all the results presented in this paper, maximization of biomass is used as the objective function value. However, the simulated annealing framework allows for the incorporation any experimental data (fluxomics, metabolomics, proteomics, etc.) and the alignment of these ecGEMs with biological reality.

**Algorithm 2** Simulated Annealing for Enzyme *k*_cat_ Optimization

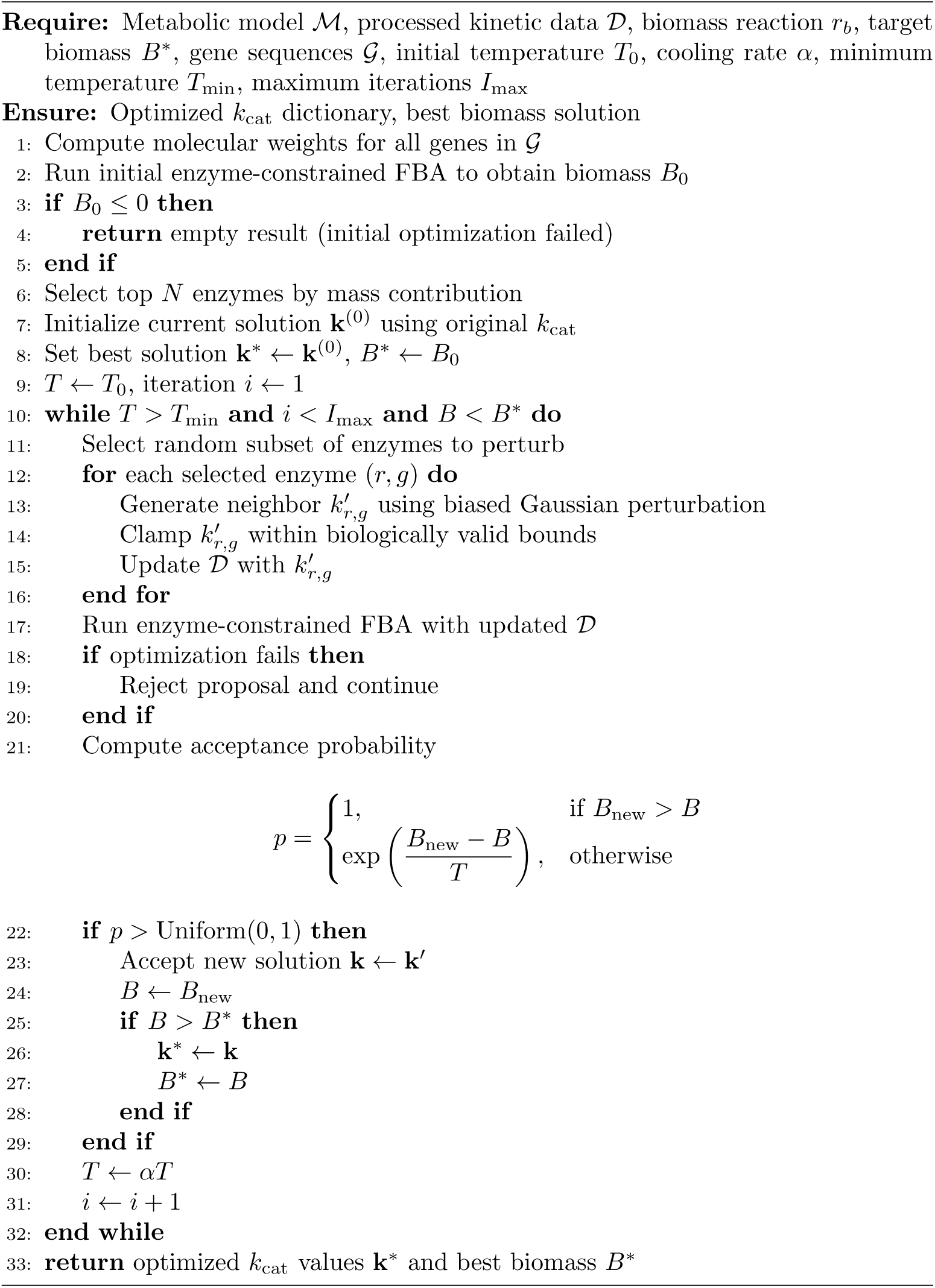

**Fig B3:**
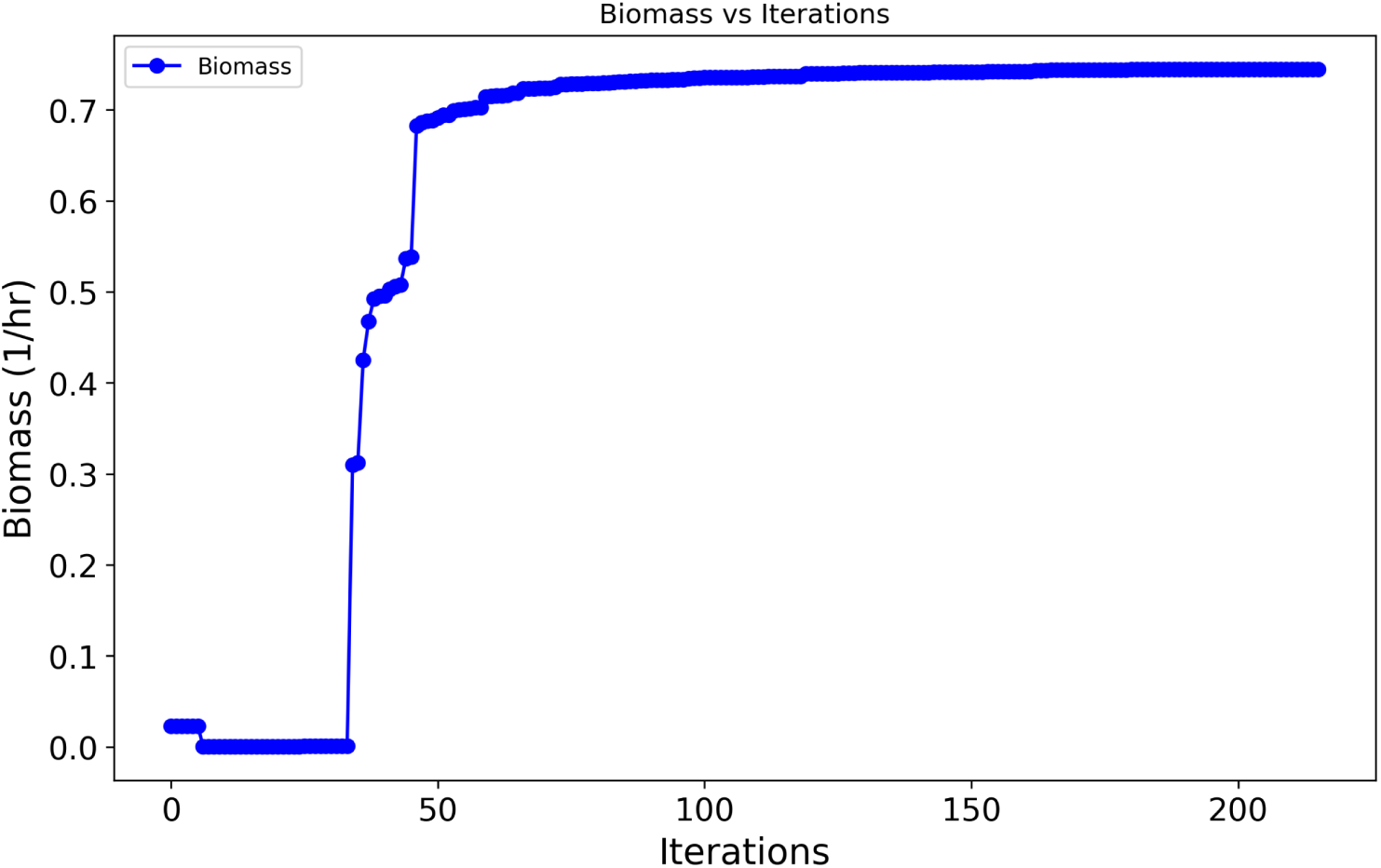
Optimization of biomass production during simulated annealing of kinetic parameters. Biomass objective value as a function of simulated annealing iterations. Early iterations exhibit low or near-zero growth, followed by a rapid transition to feasible growth states as constraints are relaxed. Subsequent iterations show gradual improvement and stabilization of biomass, indicating convergence of the optimization process toward a high-fitness kinetic parameter configuration.

**Fig B4:**
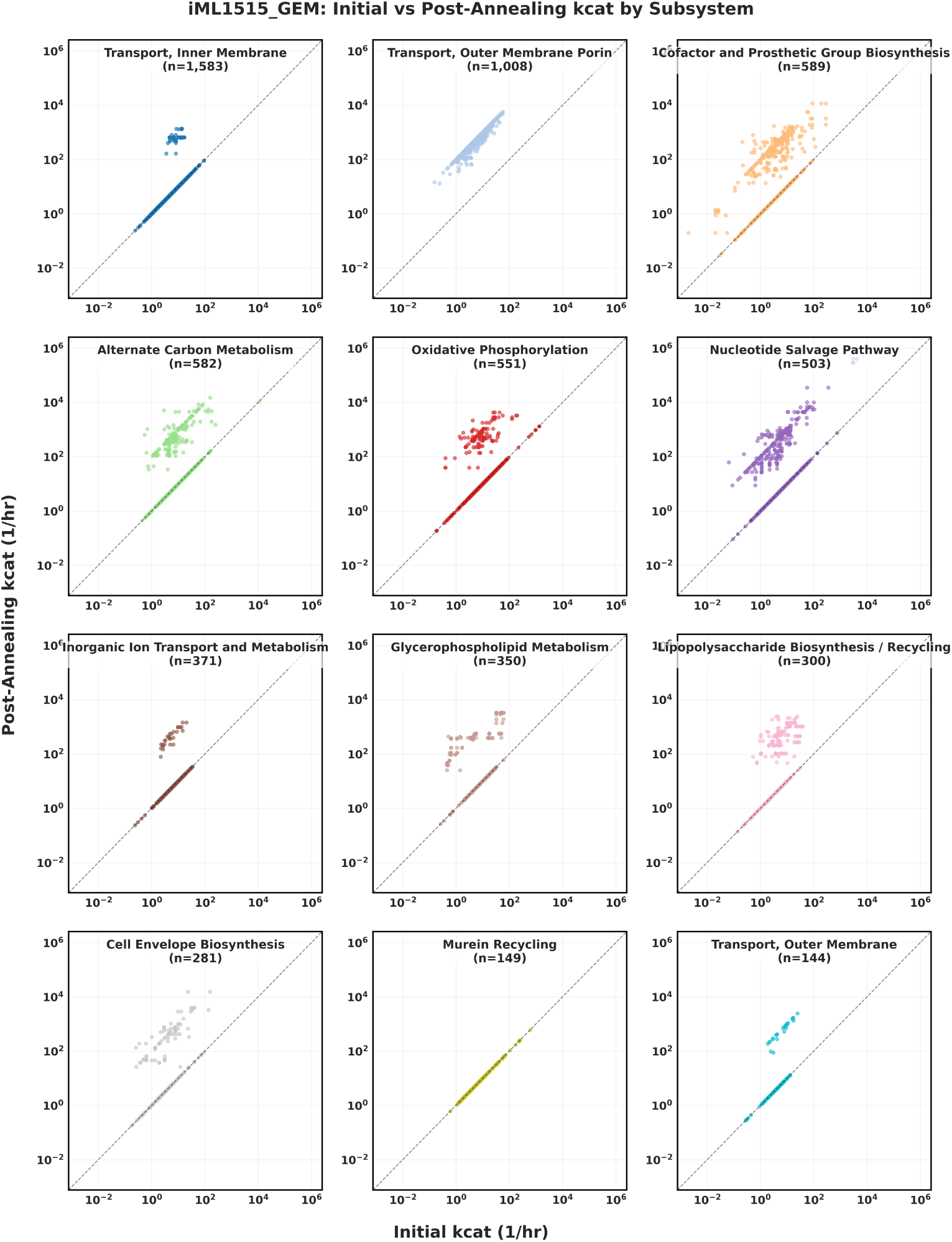
**Subsystem-specific shifts in catalytic rate constants (***k_cat_***) following simulated annealing optimization.** Scatter plots highlighting the shifts of *k_cat_* values per subsystem in *E. coli* ’s (iML1515) metabolism. Outer membrane porin transport reactions reveal consistent upward bias in the simulated annealing process. Note that there about 5,000 data points with unknown or other smaller subsystems that are excluded from this figure.

**Fig B5:**
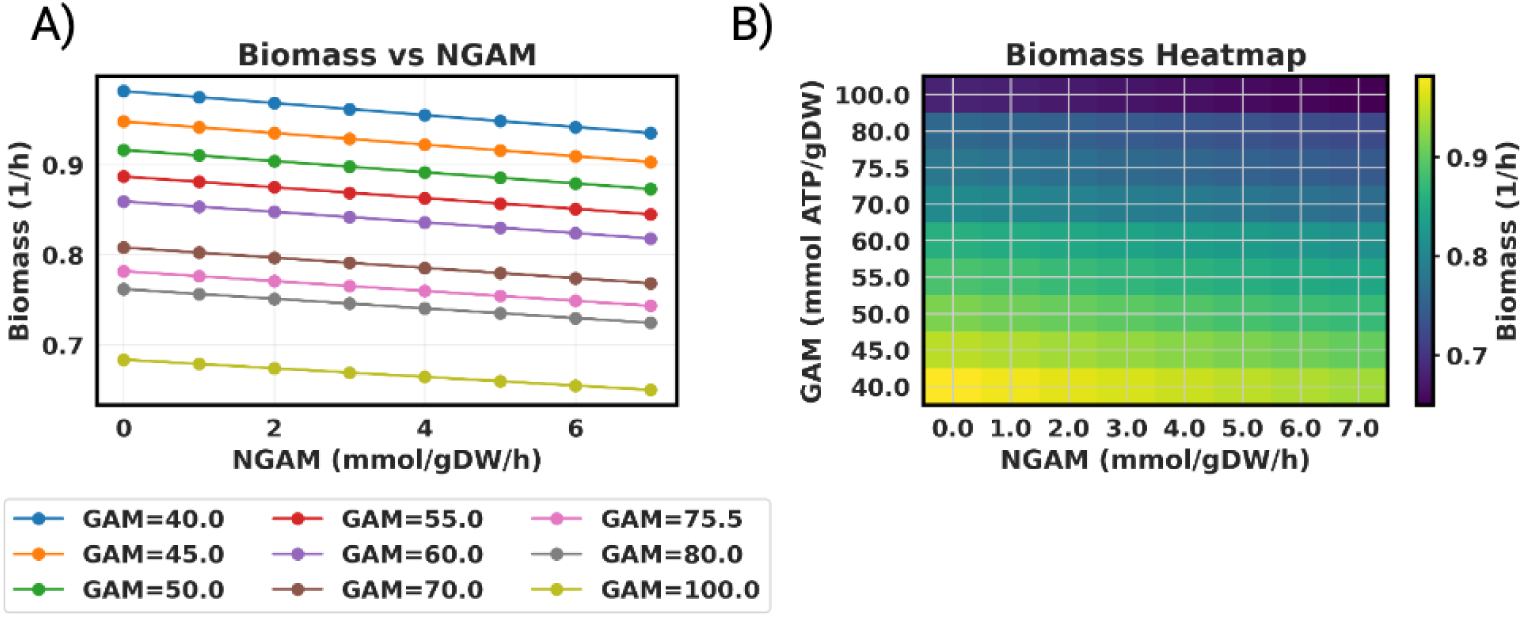
Sensitivity of predicted growth rate to growth-associated (GAM) and non-growth-associated (NGAM) maintenance parameters in *E. coli* iML1515. (A) Biomass production rate as a function of NGAM across different fixed GAM values. Increasing NGAM leads to a monotonic decrease in biomass for all GAM settings, while higher GAM values uniformly shift biomass predictions downward, indicating increased energetic cost of growth. (B) Heatmap summarizing biomass production across the GAM–NGAM parameter space. Biomass is highest in regions with low GAM and NGAM and decreases smoothly as maintenance energy requirements increase, illustrating the coupled impact of growth and maintenance energy demands on predicted fitness.

**Fig B6:**
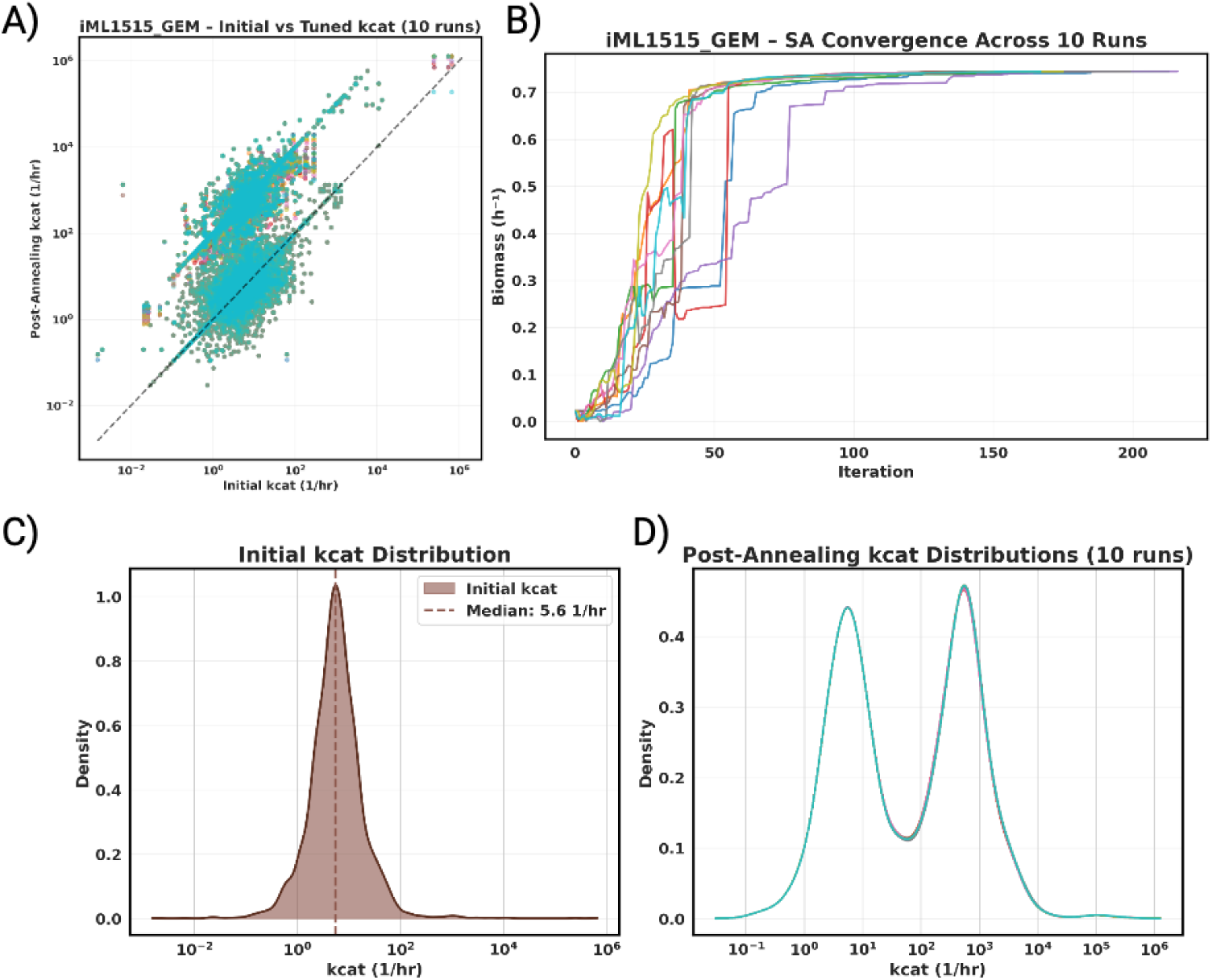
Reproducibility and convergence of simulated annealing optimization across 10 independent runs in *E. coli* iML1515. (A) Scatter plot comparing initial and post-annealing *k_cat_* values across all 10 runs. The dashed line indicates no change (*y* = *x*). The majority of points lie above the diagonal, indicating a consistent upward shift in catalytic rates across seeds, with high reproducibility in the direction and magnitude of perturbations. **(B)** Biomass production as a function of simulated annealing iteration across all 10 runs. Despite different random seeds, all runs converge to similar biomass values ( 0.75 *h^−^*^1^) within approximately 50-100 iterations, demonstrating robust convergence behavior independent of initialization. **(C)** Initial *k_cat_* distribution prior to simulated annealing, showing a unimodal distribution with a median of 5.6 *h^−^*^1^ concentrated at low turnover rates. **(D)** Post-annealing *k_cat_* distributions across all 10 runs (overlaid). All runs produce nearly identical bimodal distributions with a rightward shift relative to the initial distribution, indicating that the optimization consistently identifies similar regions of the kinetic parameter space regardless of random seed.

### B.1 Sparsity Sweep

As part of the kinGEMs pipeline design, we additionally evaluated how the size of the enzyme perturbation pool (the number of top-by-mass genes simulated annealing is eligible to perturb per iteration (*n top enzymes*)) affects feasibility, proteomics agreement, and interpretability. We compared *n top enzymes* ∈ {5, 10, 20, 50, 500} (CPI-Pred initialization, 8 growth conditions × 4 replicates; Fig B7), spanning a sparse (GECKO/ECMpy-style) regime to the broad (DLKcat-style) regime used throughout the primary results in this study.

**Fig B7:**
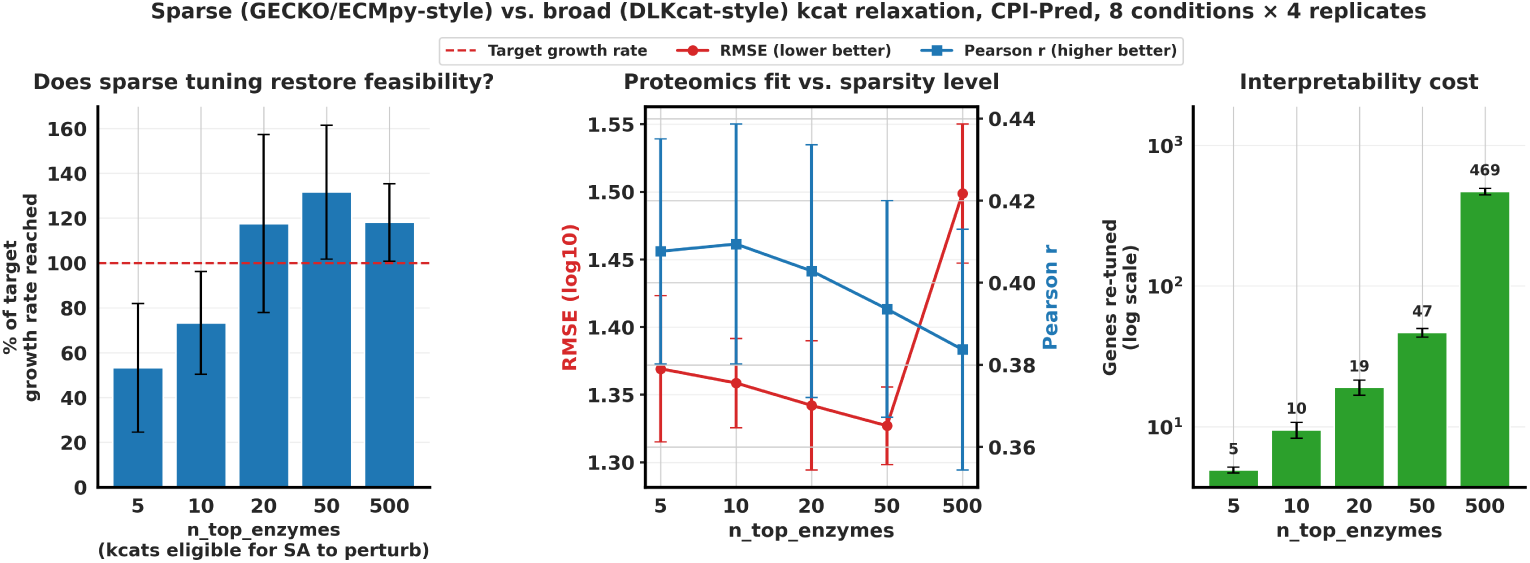
Sparse vs. broad. *k_cat_* **relaxation: effect of perturbation pool size on feasibility, proteomics fit, and interpretability.** Comparison of restricting simulated annealing to perturb only the top-*n* enzymes by mass contribution (*n* ∈ {5, 10, 20, 50, 500}), using CPI-Pred-initialized *k_cat_s* across 8 growth conditions with 4 replicates each. **Left:** Percentage of the target growth rate reached after tuning (dashed line = 100%); very small pools (*n* = 5, *n* = 10) fail to reliably restore feasible growth, while *n* ≥ 20 consistently meets or exceeds the target. **Center:** Proteomics fit (RMSE, red; Pearson *r*, blue) against Schmidt et al. proteomics, as a function of pool size. RMSE improves monotonically from *n* = 5 to *n* = 50, then degrades sharply at *n* = 500 (the broad pool used elsewhere in this study); Pearson *r* shows a milder, comparable trend. **Right:** Number of genes actually re-tuned (log scale) at each pool size, illustrating the interpretability cost of broader relaxation. Together, these results indicate that a moderate perturbation budget (*n* ≈ 20-50) achieves comparable or better feasibility and proteomics accuracy than the broad (*n* = 500) pool used for the primary results in this study, at substantially lower interpretability cost.

## Appendix C MFA Experimental Comparison

**Fig C8:**
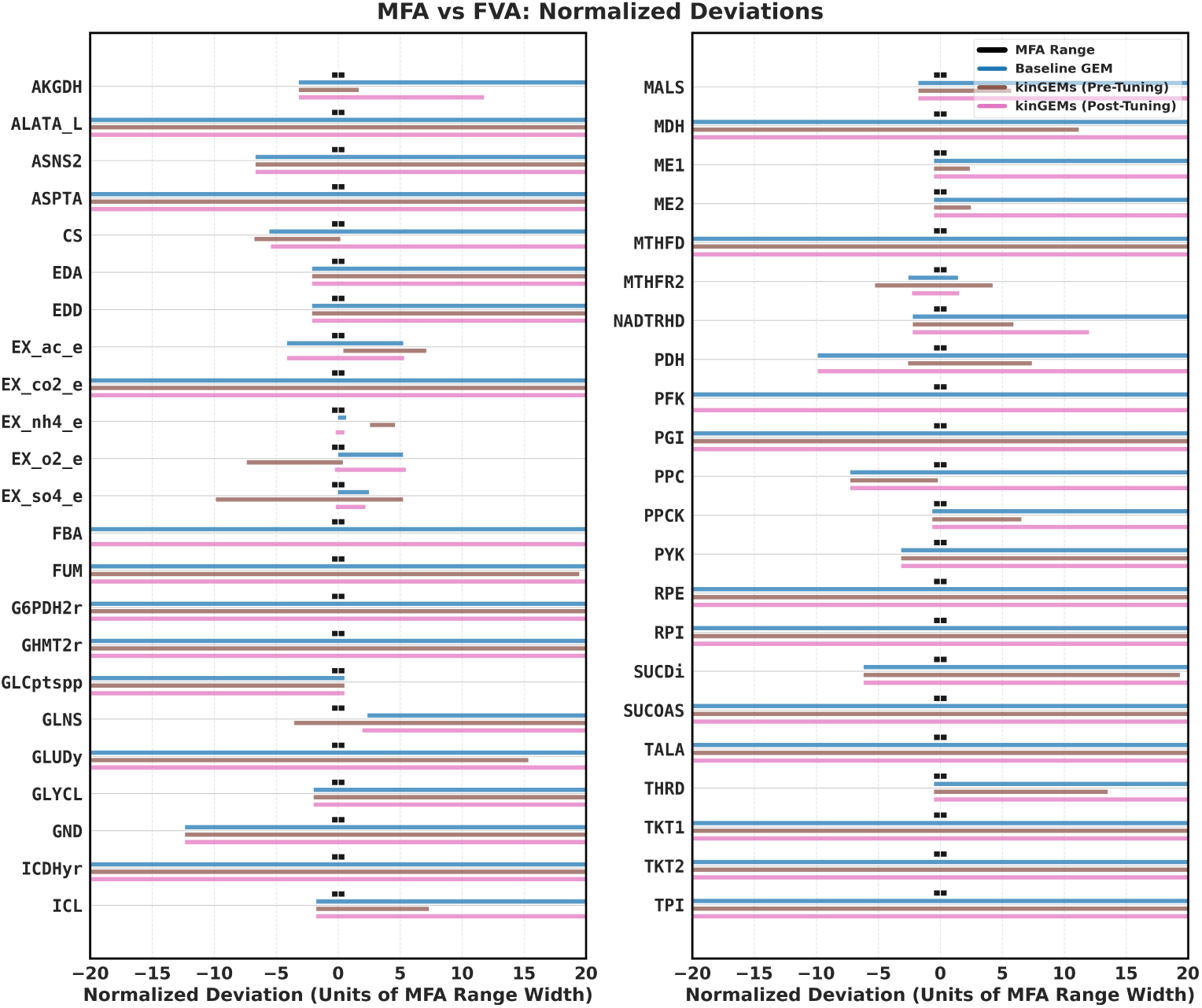
FVA Results Normalized Against Experimental MFA Values. Comparison of experimentally measured ^13^C MFA flux ranges (black) with model-predicted feasible flux ranges obtained via FVA for all available reactions. Colored horizontal bars indicate FVA ranges for the baseline GEM (blue), kinGEMs before kinetic tuning (brown), and kinGEMs after kinetic tuning (pink). Enzyme constraints substantially narrow feasible flux ranges relative to the baseline model, bringing predictions closer to experimentally observed values. Some reactions have missing kinGEMs (Pre-Tuning) FVA ranges because the optimization failed to find solutions for the reactions at that formulation.

## Appendix D Supplementary Proteomics and Phenotype Validation

This appendix provides supplementary detail supporting the proteomics and phenotypic accuracy analyses presented in Section 3.2, using the absolute quantitative proteomics dataset of Schmidt et al. [30] across 15 growth conditions and 4 *k_cat_*initialization sources (CPI-Pred, an independent deep learning predictor CatPred, literature-derived BRENDA values, and randomly sampled log-normal values).

**Table C1:**
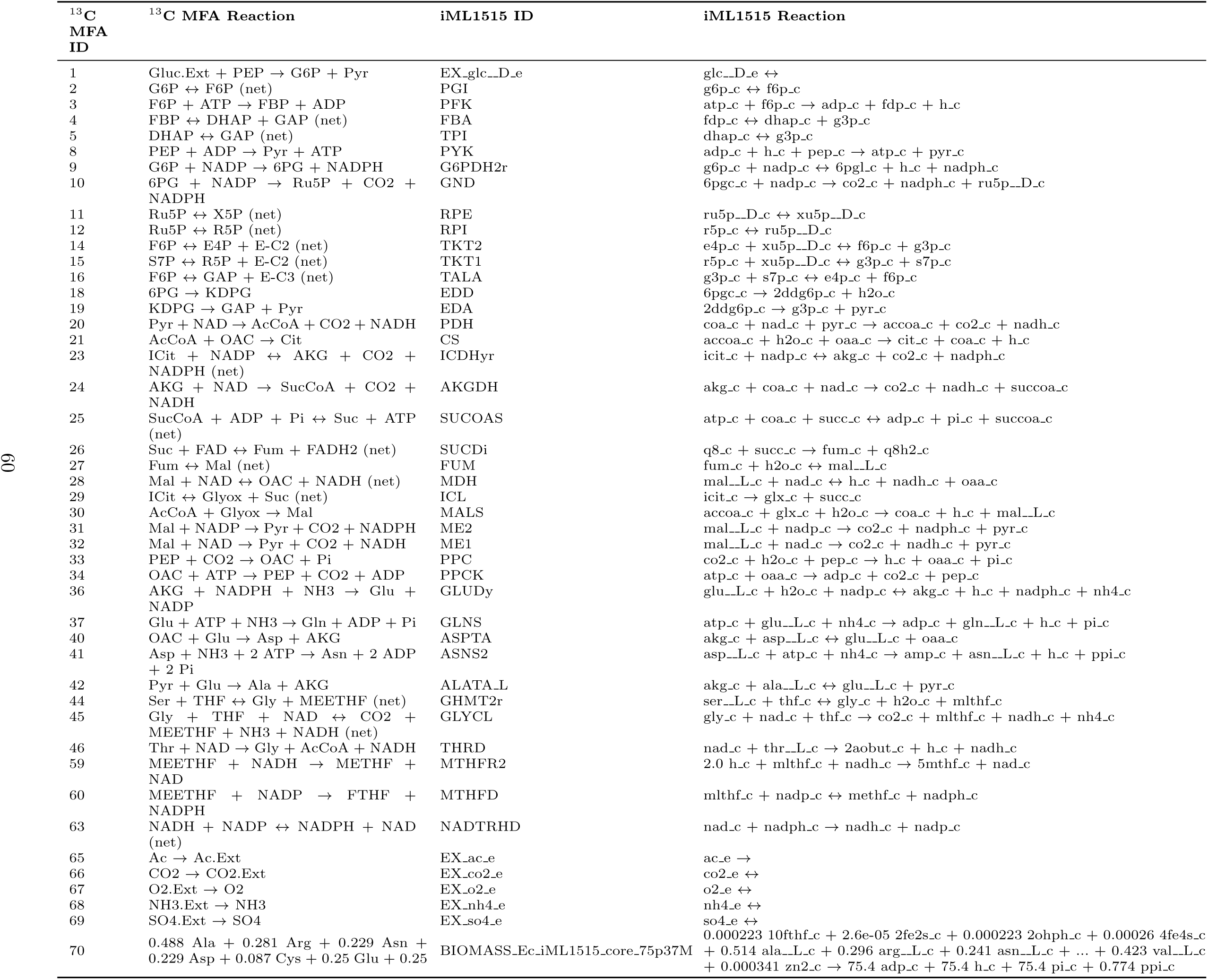
Mapping between ^13^C MFA for intracellular and exchange reactions [29] and iML1515 model reactions.

**Table C2:** Wilcoxon signed-rank tests comparing Baseline GEM against kinGEMs (post-tuning) and kinGEMs (pre-tuning) for mean-to-mean flux distance and FVA interval width (log-scale). Using glucose-normalized fluxes (rescaled to glucose uptake = 100). Effect size is reported as rank-biserial correlation *r*; *^∗^p <* 0.001.

| Metric | Comparison | Median<br>(Baseline) | Median<br>(model) | $n$ | $p$-value | $r$ |
| --- | --- | --- | --- | --- | --- | --- |
| Mean-to-mean<br>distance | vs. kinGEMs (post-tuning) | 48.54 | 41.59 | 46 | $2.77 \times 10^{-4*}$ | 0.929 |
| | vs. kinGEMs (pre-tuning) | 48.54 | 14.19 | 46 | $2.06 \times 10^{-4*}$ | 0.836 |
| FVA interval<br>width (log-scale) | vs. kinGEMs (post-tuning) | 380.23 | 292.93 | 45 | $4.33 \times 10^{-4*}$ | 0.813 |
| | vs. kinGEMs (pre-tuning) | 380.23 | 104.97 | 45 | $1.44 \times 10^{-6*}$ | 0.921 |

### D.1 Growth-rate phenotype agreement

As an additional phenotype-level check, we compared model-predicted growth rate (the FBA objective) against experimentally reported growth rates from Schmidt et al. across all 15 conditions and 4 *k_cat_*sources (n=55; Fig D9). Predicted and experimental growth rates showed strong agreement (r = 0.73, MAE = 0.12 h*^−^*^1^). We note this agreement is expected to be strong rather than an independent validation, since simulated annealing actively searches for *k_cat_* values that hit the experimental growth rate as its tuning target for each condition; it confirms that the enzyme-mass budget used throughout this study is large enough to reach physiologically realistic growth rates across all 15 conditions tested without hitting an infeasible ceiling, and is presented here for completeness rather than as free out-of-sample evidence of predictive accuracy.

**Fig D9:**
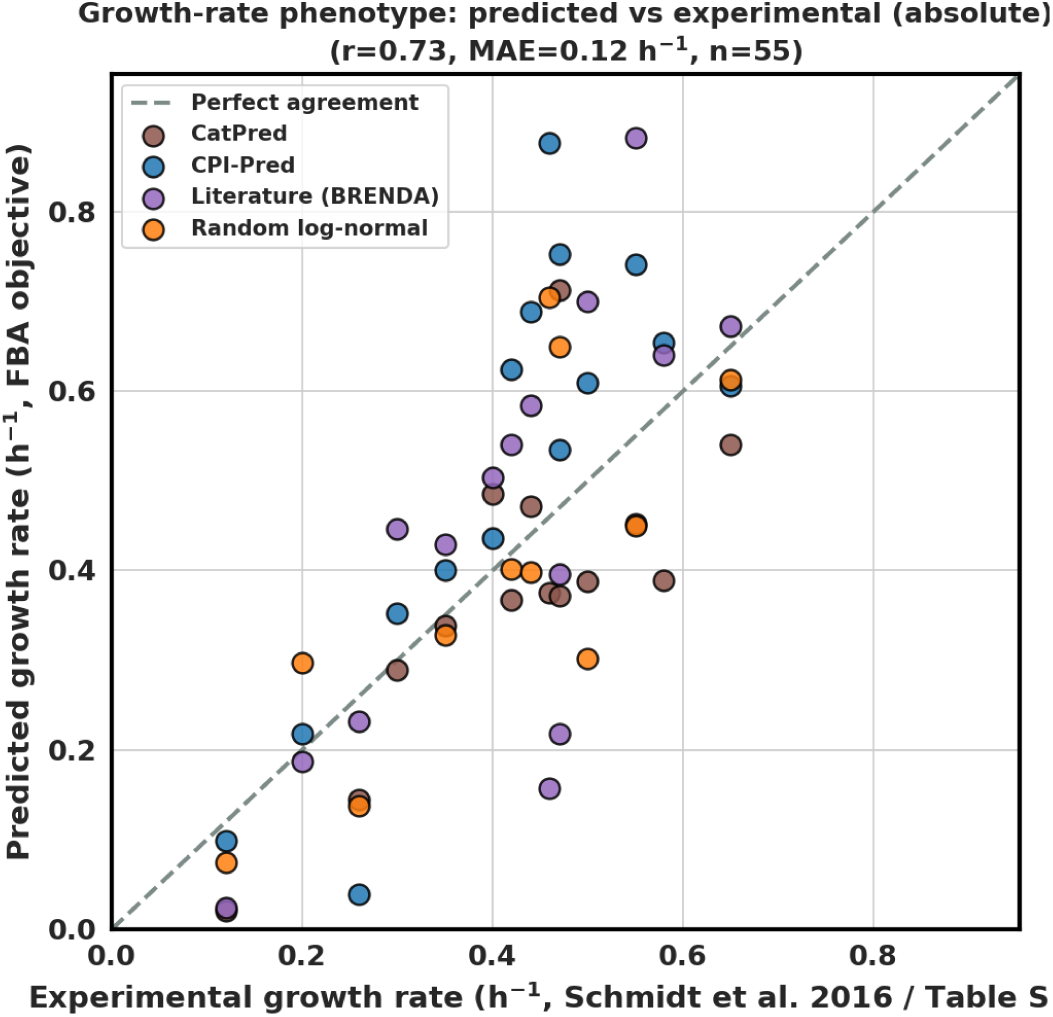
Predicted vs. experimental growth rate across 15 growth conditions and 4. *k_cat_* **sources.** Model-predicted growth rate (FBA objective, *y*-axis) vs. experimentally reported growth rate (Schmidt et al. 2016, Table S23, *x*-axis) for all 15 growth conditions and 4 *k_cat_*sources (*r* = 0.73, MAE = 0.12 h*^−^*^1^, *n* = 55). See Section D.1 for discussion of why this agreement should not be read as independent validation.

### D.2 Generalization and Specificity of Simulated-Annealing-Tuned Kinetic Parameters

A central question for any parameter-tuning procedure is whether the tuned parameters capture generalizable, biologically meaningful structure or simply overfit to the specific objective and data used during optimization. We assessed this along two independent axes: whether a *k_cat_* vector tuned under one growth condition generalizes to a different condition’s medium (Section D.2.2), and whether proteomics-aware tuning targeted at one set of genes generalizes to genes in other, untouched metabolic pathways within the same condition (Section D.2.1). These two checks yield different answers: growth-rate-driven *k_cat_* tuning transfers well across growth conditions, while gene-subset-targeted proteomics-aware tuning does not demonstrate reliable transfer to a held-out pathway.

#### D.2.1 Pathway-level specificity of proteomics-aware tuning

Genes were split by metabolic pathway into a training set (energy metabolism; glycolysis, TCA cycle, pentose phosphate pathway, oxidative phosphorylation, and pyruvate metabolism; the objective genes for the proteomics-aware tuning phase) and a heldout test set (ten amino acid biosynthesis pathways, never touched by proteomics-aware tuning). For each of the 15 growth conditions and 4 *k_cat_* sources, we compared RMSE against Schmidt et al. proteomics before and after proteomics-aware tuning, separately for genes in the training pathway and genes in the held-out pathway (Fig D10).

**Fig D10:**
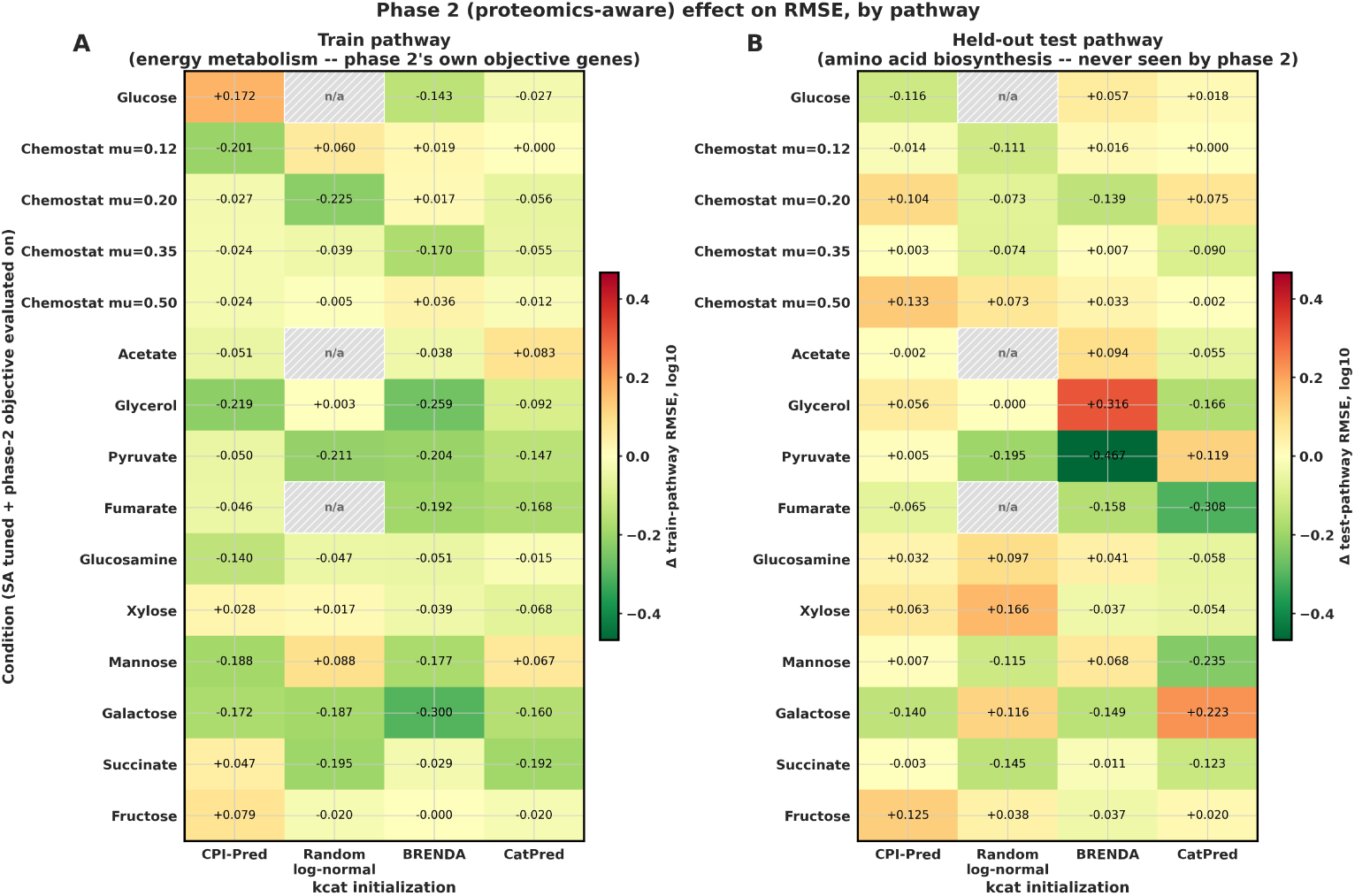
Proteomics-aware tuning generalizes from trained to held-out metabolic pathways. Change in RMSE against Schmidt et al. proteomics following proteomics-aware (Phase 2) tuning, relative to growth-rate-only (Phase 1) tuning, by growth condition (rows) and *k_cat_* initialization source (columns). **(A)** Change in RMSE for the training pathway (energy metabolism: glycolysis, TCA cycle, pentose phosphate pathway, oxidative phosphorylation, pyruvate metabolism), the direct objective of Phase 2 tuning. **(B)** Change in RMSE for the held-out test pathway (amino acid biosynthesis, 10 pathways), never included in the Phase 2 objective. Green indicates improved (lower) RMSE after proteomics-aware tuning; orange/red indicates worsened RMSE. Hatched cells (n/a) indicate condition-source combinations for which simulated annealing did not converge to a feasible solution. Panel B’s condition-source RMSE changes were not statistically significant in any completed run (95% bootstrap confidence intervals; Methods 5.6.3), and should not be interpreted as evidence that proteomics-aware tuning generalizes to genes outside its direct training objective.

Proteomics-aware tuning reduced RMSE for the training (energy metabolism) pathway in the majority of condition-source combinations (green cells, panel A), as expected given these genes were the direct tuning objective and only proteomics-fit-improving moves are accepted by construction (Methods 5.6.3). The held-out amino acid biosynthesis pathway (panel B) shows a visually mixed pattern of improvement and degradation across condition-source combinations; however, this change was not statistically significant in any completed run when assessed via 95% bootstrap confidence intervals (percentile method, resampling matched gene pairs; Methods 5.6.3). We therefore do not interpret panel B as evidence that proteomics-aware tuning generalizes to genes outside its direct training objective; the point-estimate RMSE changes shown here should be read alongside this caveat rather than as a standalone indicator of transfer. Cells marked n/a correspond to condition-source combinations for which the underlying simulated annealing run did not converge to a feasible solution.

#### D.2.2 Condition-level generalization: cross-condition transfer

We additionally tested whether a *k_cat_* vector tuned under one growth condition retains its proteomics fit when applied, without further tuning, to a different condition’s medium. For each of the 15 conditions and 4 *k_cat_* sources, we computed the native (matched-condition) RMSE against Schmidt et al. proteomics, expressed relative to the grand mean RMSE across all condition-source cells (1.541; panel A), and the cross-condition transfer effect, defined as the mean RMSE across all other target conditions minus the native RMSE (panel B; Fig D11).

**Fig D11:**
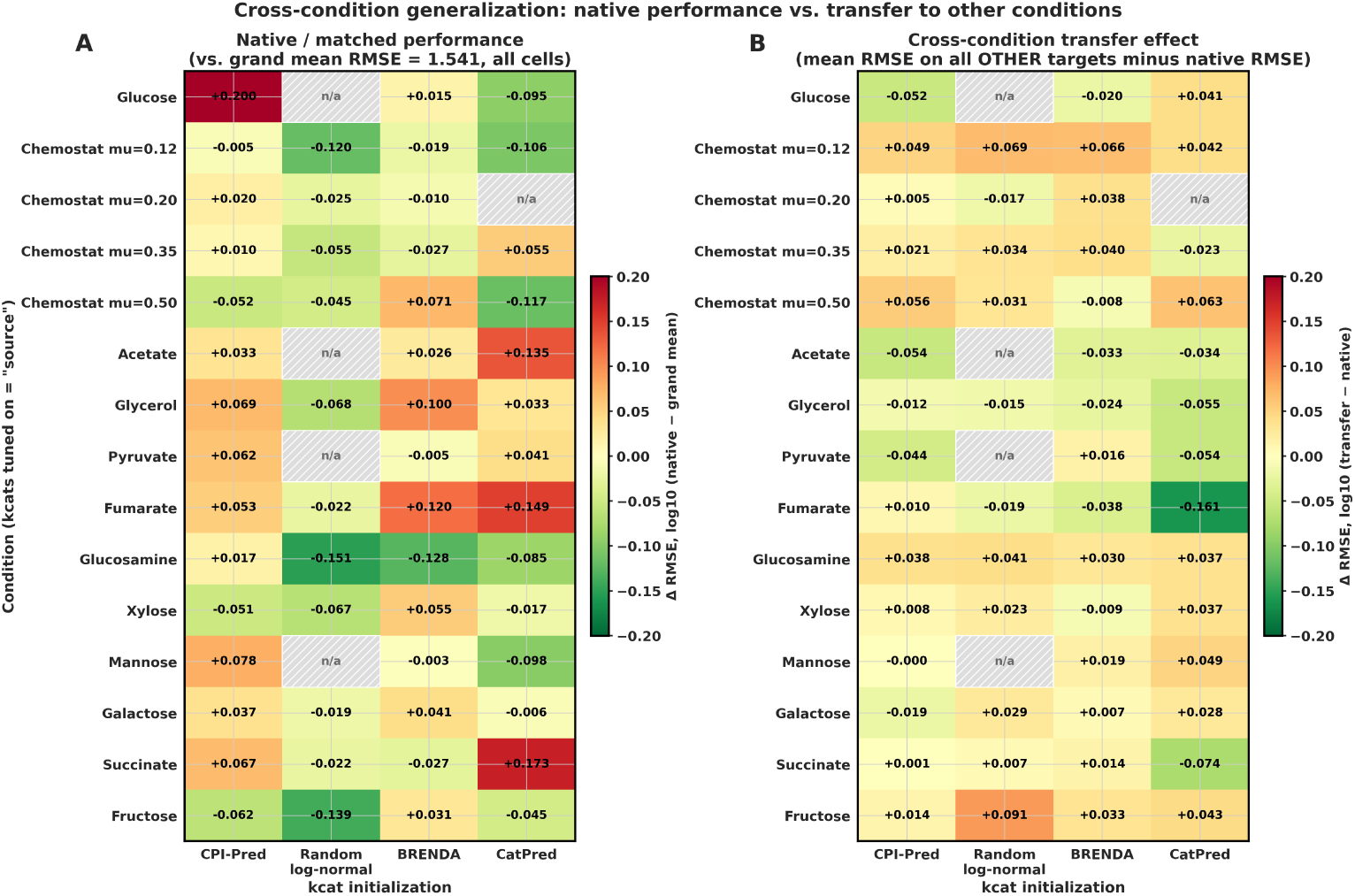
Cross-condition transfer of simulated-annealing-tuned. *k_cat_***vectors is comparable to native performance. (A)** Native (matched-condition) RMSE against Schmidt et al. proteomics, expressed as a deviation from the grand mean RMSE across all 15 conditions × 4 *k_cat_*sources (grand mean = 1.541, log10 scale). Red indicates native RMSE worse than the grand mean; green indicates better than the grand mean. **(B)** Cross-condition transfer effect: mean RMSE when a condition’s tuned *k_cat_* vector is applied to all other target conditions’ media, minus that condition’s own native RMSE. Values near zero (pale cells) indicate transfer performance comparable to native performance; red indicates worse transfer (unfavorable), green indicates better transfer (favorable) than native. Hatched cells (n/a) indicate condition-source combinations excluded due to non-convergent simulated annealing runs. The small magnitude and inconsistent sign of panel B across conditions and sources indicates no systematic overfitting of tuned *k_cat_* vectors to their training condition.

Consistent with the main-text summary statistics (mean native *r* = 0.336 vs. mean cross-condition *r* = 0.335; Section 3.2), the transfer effect in panel B is small and inconsistently signed across conditions and *k_cat_* sources (values clustering near zero, with roughly comparable numbers of positive and negative deltas), indicating no systematic degradation when *k_cat_* vectors are applied outside their original tuning condition. A small number of condition-source pairs show larger transfer penalties (e.g., Fumarate with CatPred, Δ = −0.161 in the favorable direction; several Chemostat conditions show modest positive (i.e., unfavorable) transfer deltas in the +0.03 to +0.07 range), but no *k_cat_* source or condition shows a consistent, large-magnitude generalization failure. Cells marked n/a correspond to condition-source combinations excluded due to non-convergent simulated annealing runs.

### D.3 Condition constraints for *Rhodotorula toruloides* case study

Substrate uptake and O_2_ consumption were imposed as maximum rates; CO_2_ and reported by-products were allowed up to the measured secretion rates. On xylose, xylitol and D-arabinitol secretion were bounded by the measured rates; on acetate, citrate secretion was also bounded. Glucose, xylose, and acetate exchanges that were not the intended carbon source were blocked.

**Table D3:** Constraints for the *R. toruloides* reconstructions. Units: *µ*_goal_ in h*^−^*^1^; *E*_tot_ and biomass fractions in g gDW*^−^*^1^; uptake in mmol gDW*^−^*^1^ h*^−^*^1^. Biomass

| Cond. | Carbon / N | $\mu_{goal}$ | $E_{tot}$ | Biomass | | | |
| --- | --- | --- | --- | --- | --- | --- | --- |
|  |  |  |  | Prot. | Lip. | Carb. | Uptake |
| Gexp | glc / urea | 0.191 | 0.08 | 0.64 | 0.05 | 0.20 | 2.489 |
| GNlim | glc / urea | 0.021 | 0.04 | 0.23 | 0.48 | 0.20 | 0.41 |
| Xexp | xyl / $NH_4^+$ | 0.054 | 0.07 | 0.44 | 0.20 | 0.30 | 1.86 |
| XNlim | xyl / $NH_4^+$ | 0.019 | 0.03 | 0.20 | 0.28 | 0.46 | 0.435 |
| Aexp | ac / $NH_4^+$ | 0.073 | 0.09 | 0.39 | 0.18 | 0.37 | 6.1 |
| ANlim | ac / $NH_4^+$ | 0.012 | 0.05 | 0.22 | 0.34 | 0.28 | 1.971 |

**Table D4:** Exchange bounds for the *R. toruloides* reconstructions.

| Cond. | $O_2$ | $CO_2$ | By-products | | | | |
| --- | --- | --- | --- | --- | --- | --- | --- |
|  |  |  | N | glyc | xylitol | ara | cit |
| Gexp | -5.6 | 7.3 | urea | 0.049 | — | — | — |
| GNlim | -1.16 | 1.440 | urea | -0.007 | — | — | — |
| Xexp | -2.678 | 2.845 | $NH_4^+$ | 0 | 0.223 | 0.367 | — |
| XNlim | -1.028 | 1.277 | $NH_4^+$ | 0 | 0.004 | 0.077 | — |
| Aexp | -7.144 | 5.224 | $NH_4^+$ | 0 | — | — | 0.122 |
| ANlim | -2.265 | 2.229 | $NH_4^+$ | 0 | — | — | -0.043 |

## Appendix E Running kinGEMs on the BiGG Database

**Table E5:**
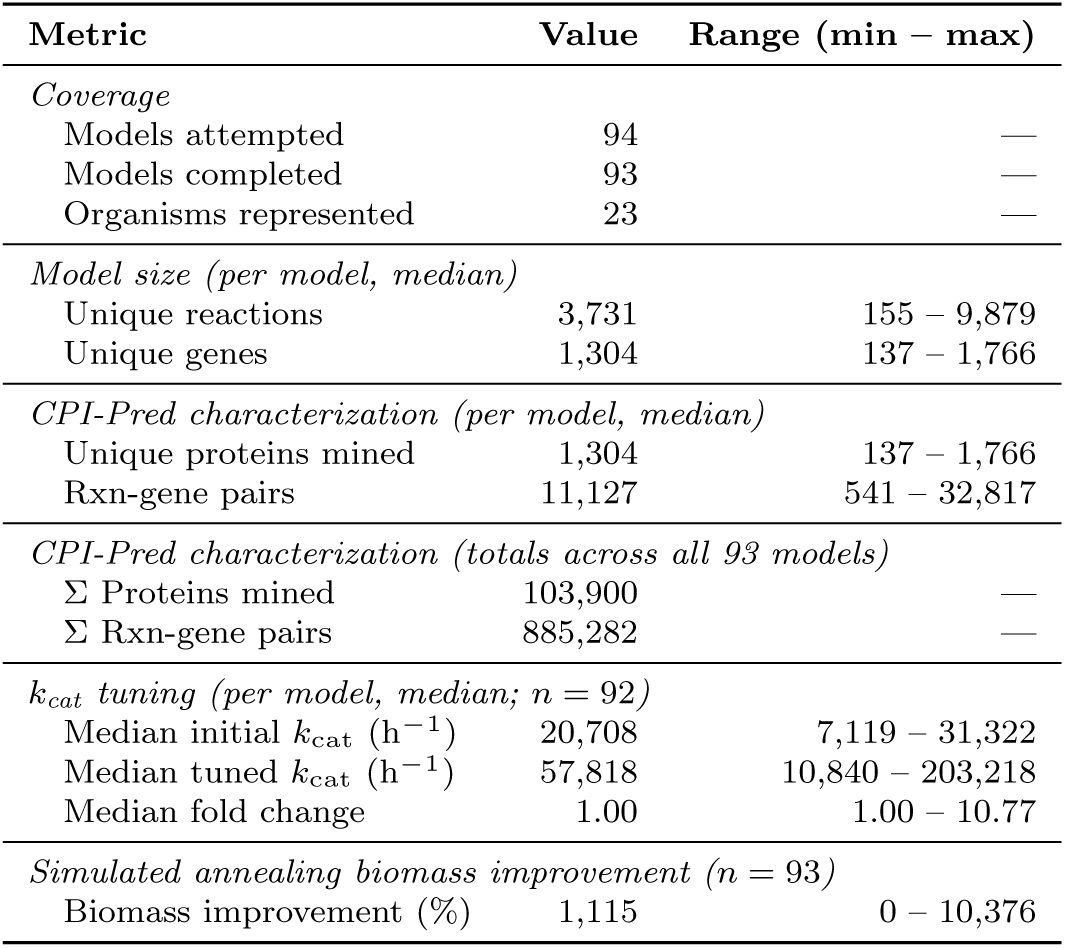
Aggregate statistics for the kinGEMs pipeline applied across 93 completed BiGG genome-scale metabolic models spanning 23 organisms.

**Fig E12:**
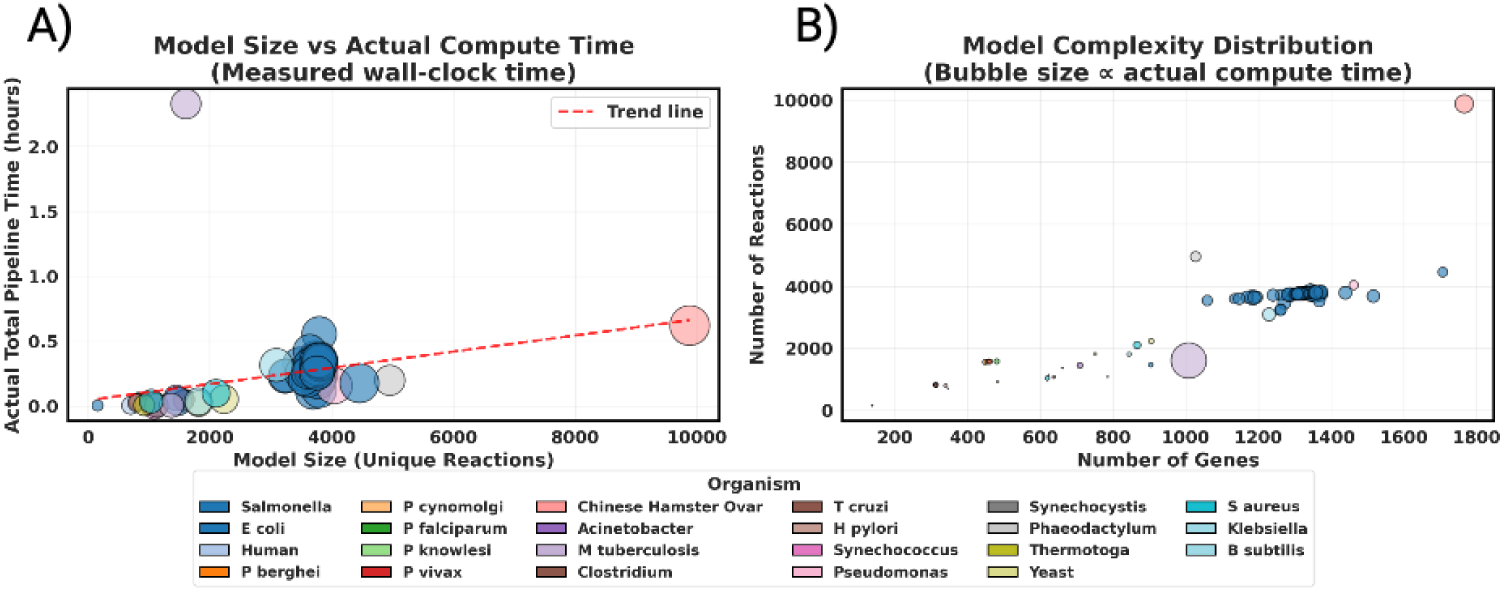
Computational scalability of the kinGEMs pipeline across 93 BiGG genome-scale metabolic models spanning 23 organisms. **(A)** Actual total pipeline wall-clock time as a function of model size (unique reactions) across all processed models. The dashed red trend line indicates a approximately linear scaling relationship between model size and compute time, with the majority of models completing in under 0.5 hours. The Chinese Hamster Ovary model, the largest model processed ( 9,800 reactions), required approximately 0.6 hours, demonstrating tractable runtime even for large-scale models. Bubble size is proportional to actual compute time. **(B)** Model complexity distribution across all processed organisms, with each bubble representing a model positioned by its number of genes (x-axis) and number of reactions (y-axis), and bubble size proportional to actual compute time. Models span a broad range of biological complexity, from small prokaryotic models (135 reactions) to large eukaryotic models (9,800 reactions), covering diverse taxa including bacteria, parasites, fungi, cyanobacteria, and mammalian cell lines.

**Table E6:**
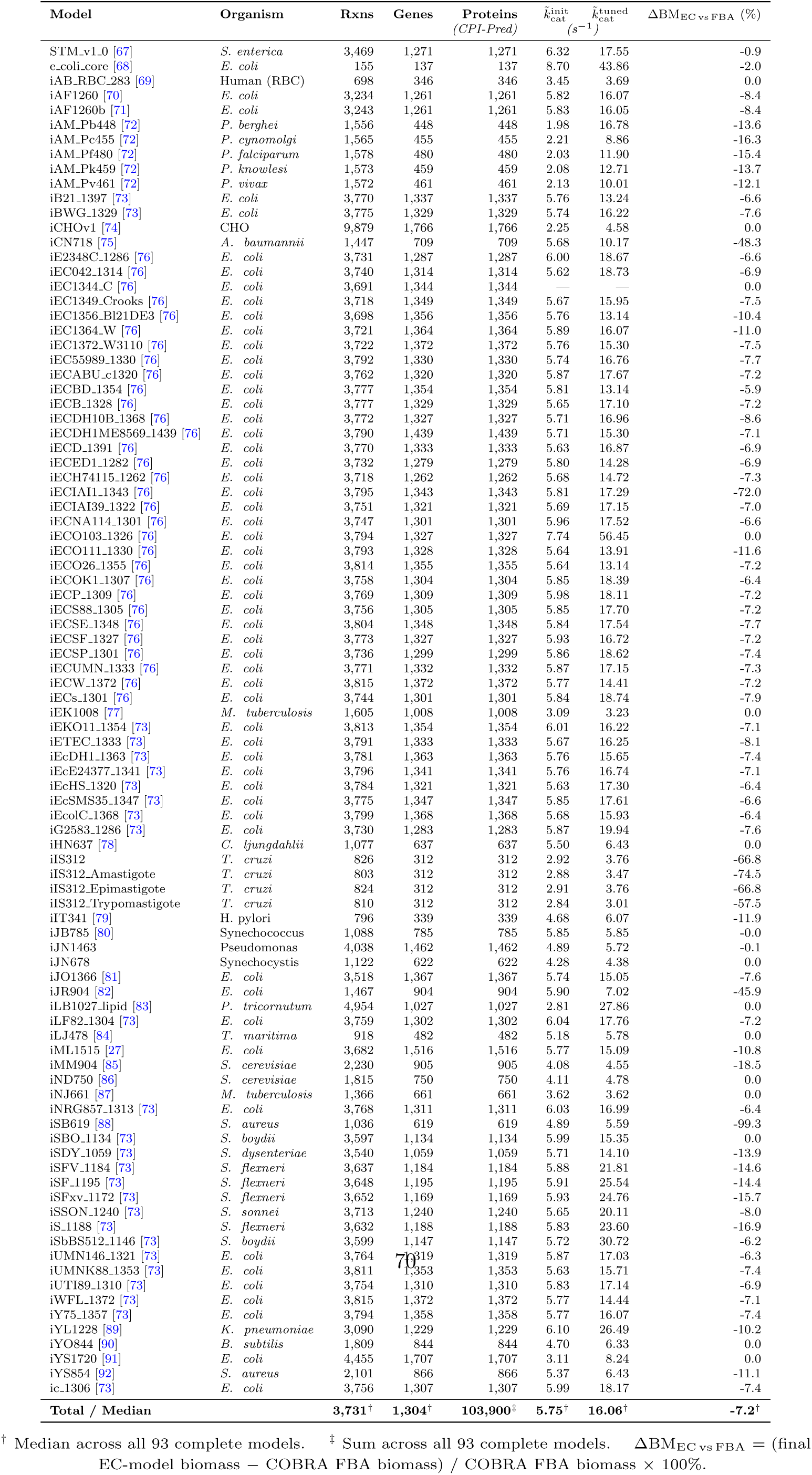
Summary statistics across genome-scale metabolic models (*n* = 93 complete runs out of 94 attempted).

## Notes

https://github.com/LMSE/kinGEMs

### Competing Interest Statement

The authors have declared no competing interest.

### Summary of Updates

This version of the manuscript to add some additional analyses on non-E. coli organisms as well as some overflow metabolism analyses.

